# Attenuated interferon signalling in alveolar epithelium limits resistance to *Streptococcus pyogenes*

**DOI:** 10.64898/2026.04.15.718576

**Authors:** Jeremy Wiyana, Declan Turner, Sahel Amoozadeh, Pooja Venkat, Katelyn Patatsos, Hannah Frost, Josh Osowicki, Jasmyn Voss, Kaneka Chheng, Kristy Azzopardi, Natalie Caltabiano, Mark Davies, Mirana Ramialison, Catherine Satzke, Fernando J Rossello, Andrew Steer, Ed Stanley, Rhiannon B. Werder

## Abstract

The upper respiratory tract is a primary niche for *Streptococcus pyogenes* colonisation and disease. Lower respiratory tract infection (pneumonia) is the most common invasive *S. pyogenes* syndrome. Studies have not previously examined how epithelial cells, from the airway to the alveolus, respond to *S. pyogenes* infection. Here, we established a scalable human *in vitro* model by differentiating induced pluripotent stem cells (iPSCs) into mature pseudostratified airway epithelium or alveolar type 2 epithelial cells, cultured at air-liquid interface and infected with *S. pyogenes* (M1_UK_ and M75 strains). Both strains attached to iPSC-derived lung epithelial cells, with significantly greater adherence to the airway epithelium by M75 compared to M1_UK_. Moreover, invasion by both *S. pyogenes* strains of alveolar epithelial cells was greater than for the airway epithelium. Dynamic *S. pyogenes* gene expression changes were evident between 6 and 24 hours after infection, which was influenced by the infected cell type; however, virulence genes were not significantly altered. While infection of the airway epithelium induced rapid and dynamic inflammatory signalling, the alveolar epithelium demonstrated augmented cell death and mounted a transcriptional pro-inflammatory and proliferative response that was uncoupled from cytokine secretion. The airway epithelium model exhibited consistently higher baseline type I interferon (IFN) signalling than the alveolar epithelium. Invasion by *S. pyogenes* and inflammation was significantly reduced in IFN-β-treated alveolar epithelial cells. In summary, we have established the first model of *S. pyogenes* infection in physiologically relevant airway and alveolar epithelial cells. Our findings suggest that host responses to infection are influenced by lung compartment, the *S. pyogenes* strain type, and infection timepoint, highlighting context-specific pathways that could be leveraged therapeutically.

## INTRODUCTION

Pneumonia remains the leading infectious cause of death in children worldwide, accounting for 15% of all deaths in those under the age of 5 (1). Pneumonia is characterised by excessive inflammation within the alveoli, typically caused by bacterial, viral or fungal infection. While most *Streptococcus pyogenes* (Group A Streptococcus) infections in the upper respiratory tract are asymptomatic or mild, less common invasive lower respiratory tract infections can be devastating. Pneumonia is the most frequent invasive *S. pyogenes* disease syndrome in children (2–4) and is more severe than pneumonia caused by other common pathogens (5–7). A recent study found that 92% of affected patients were admitted to an intensive care unit (8). Globally, severe *S. pyogenes-*pneumonia is fatal in more than one third of cases, despite antibiotic therapy and supportive care (9–11). There is no vaccine to prevent these severe infections. Greater understanding of the pathogenesis of lower respiratory tract infection by *S. pyogenes* will inform vaccine development efforts and improved treatment strategies.

The severity and outcome of *S. pyogenes* infections are shaped by both bacterial and host factors. *S. pyogenes* is a highly adapted human-restricted pathogen. Certain strain types (e.g., M1) are widely prevalent, more associated with invasive infections (12–15), and exhibit remarkable capacity to adapt and spread globally (12, 16, 17). Specific bacterial genes are also enriched among invasive isolates, including antibiotic resistance elements, regulators of virulence, and superantigens (17–20). Moreover, severe invasive disease develops when bacteria gain access to normally sterile sites, including blood, soft tissues or the lung alveoli (21). While controlled human infection studies have extensively characterised immune responses in *S. pyogenes* pharyngitis (22, 23), ethical constraints preclude a comparable challenge model for *S. pyogenes* pneumonia. As a consequence, respiratory *S. pyogenes* infection studies have typically relied on immortalised cell lines (24–26), which lack the complexity needed to recapitulate epithelial responses to infection. In contrast, studies in paired primary human airway and alveolar epithelial cells are limited by the difficulty of obtaining and maintaining these cells in culture.

iPSC-derived respiratory models are increasingly being used to study respiratory infections with human-restricted pathogens. Our group and others have developed directed differentiation approaches to create airway and alveolar epithelial cells (27–31). These cells closely resemble human airway or alveolar epithelium in terms of gene expression, key cell type-specific functions (e.g., mucociliary clearance, surfactant production) (28, 29), are scalable (32), and recapitulate disease-specific phenotypes (33–35). Moreover, by incorporating an Air-Liquid Interface (ALI) cell culture format, these models mimic the native respiratory surface with correct apical and basolateral polarity. The accessible apical surface provides an optimal platform for the study of respiratory infections (e.g., SARS-CoV-2, RSV) (30, 32, 36–40). Here, we leverage iPSC-derived airway and alveolar epithelial ALI systems to explore responses to *S. pyogenes* infection. We find that the bacteria adhere to and invade epithelial cells, with strain- and cell type-specific differences occurring despite minimal shifts in bacterial virulence gene expression. By mapping the epithelial response across lung cell types, we establish a scalable human model for *S. pyogenes* respiratory infection, which highlights a key role for the epithelium in influencing the trajectory of host–pathogen interactions.

## RESULTS

### *S. pyogenes* attachment and invasion is strain- and compartment-dependent in iPSC derived lung epithelium

To assemble an *in vitro* model of respiratory *S. pyogenes* infection, we generated ALI cultures containing multiciliated, club, goblet, and basal airway epithelial cells (iAEC) or surfactant-producing type 2 alveolar epithelial cells (iAT2) (Figure 1A). In iAEC or iAT2 cultures infected with *S. pyogenes* (M1_UK_ strain, at 1×10^6^ CFU/ml or 0.5 MOI) for 6 hours we observed chains of *S. pyogenes* compared with uninfected controls (Figure 1B). *S. pyogenes* produces adhesins and invasins to facilitate attachment and intracellular invasion (41–44), and their expression can vary across strains (45). To investigate the attachment to, and/or invasion of, different cell types, iAECs or iAT2s were incubated with the *S. pyogenes* strains M1_UK_ or M75 (at 1×10^6^ colony forming units (CFU_/ml) for 6 hours. M1_UK_ is a globally prevalent strain frequently responsible for fatal invasive infections (17), whereas the M75 strain is capable of milder superficial infection such as pharyngitis (46). After this incubation, live adherent or intracellular bacteria were quantified by culture (CFU). Both strains attached to both cell types, with the M75 strain adhering significantly more to iAECs than the M1_UK_ strain (Figure 1C). Adherence of the M1_UK_ strain was higher for the iAT2s compared with the iAECs (Figure 1C). Moreover, a trend of greater invasion was observed in the iAT2s by both *S. pyogenes* strains compared to iAECs (Figure 1D). Transmission electron microscopy (TEM) similarly revealed attachment of *S. pyogenes* to iAECs and iAT2s, with rare instances of intracellular localisation following infection of iAT2s (Figure 1E-G). Of note, intracellular *S. pyogenes* displayed an altered surface morphology, consistent with shedding of the hyaluronic acid capsule. Taken together, these findings suggest that both M1_UK_ and M75 strains of *S. pyogenes* are capable of adhering to, and in rare cases invading, iPSC-derived airway and alveolar epithelial cells.

**Figure 1.**
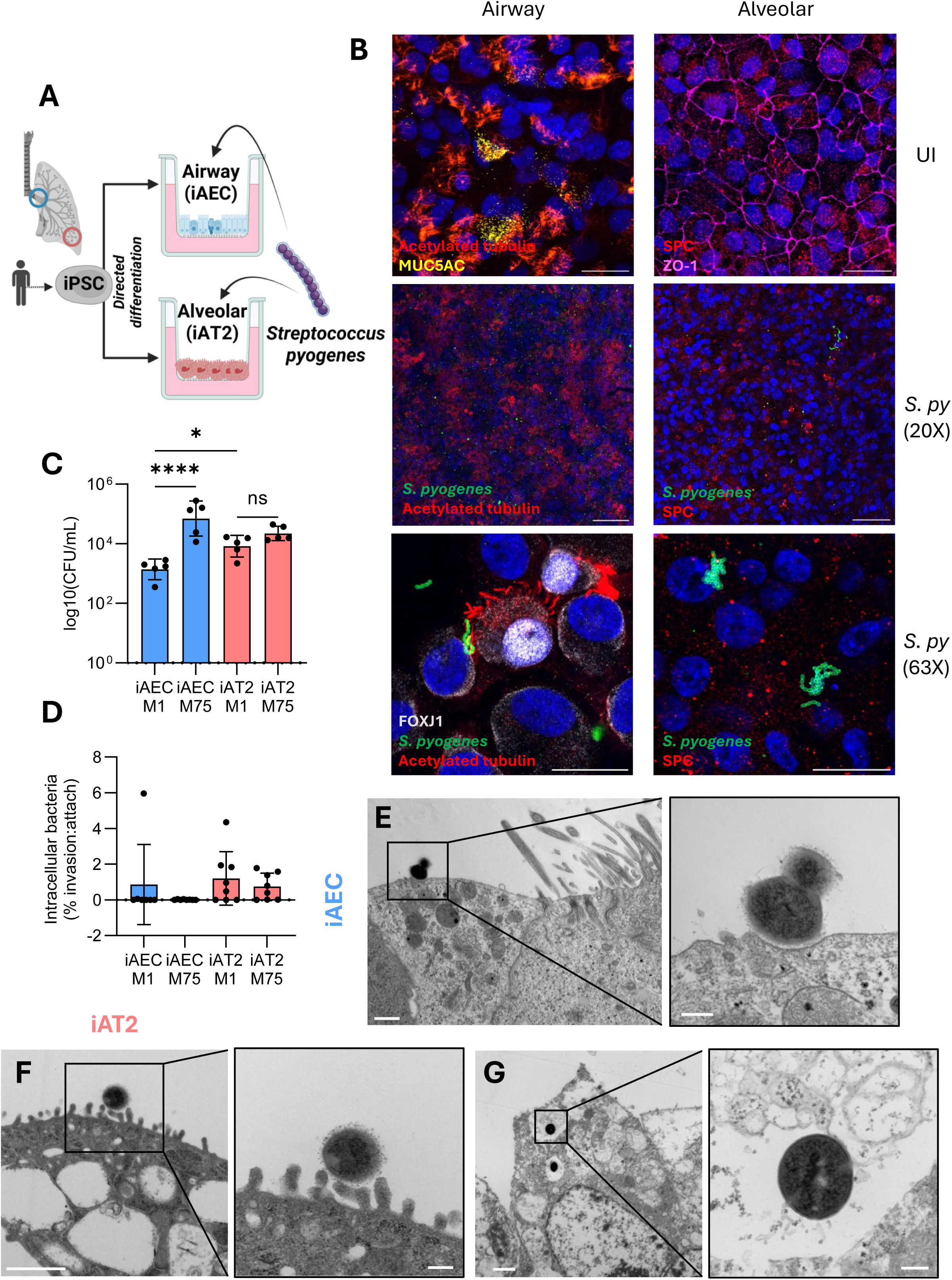
iPSC-derived airway and alveolar epithelial response to *S. pyogenes* infection. iPSC-derived airway epithelial cells (iAECs) and type 2 alveolar epithelial cells (iAT2s) were infected with *S. pyogenes* (M1_UK_ or M75, at 1×10^6^ CFU/ml) for 6 hours. **(A)** Schematic of the iPSC-derived infection model. Human iPSCs underwent directed differentiation to airway epithelial cells or iAT2s, which were further matured at air-liquid interface (ALI) prior to infection with *S. pyogenes*. **(B)** Confocal imaging of iAECs (left) stained for DAPI (blue), acetylated tubulin (red), MUC5AC (yellow), *S. pyogenes* (green) and FOXJ (white), and iAT2s (right) stained for DAPI (blue), surfactant protein C (SPC) (red), ZO-1 (pink) and *S. pyogenes* (green). Scale bar = 20µm. **(C-G)** *S. pyogenes* attaches and invades iPSC-derived airway and alveolar epithelial cells. **(C)** Live bacterial attachment and **(D)** invasion were quantified via colony forming unit (CFU) assay. n = 5-8 experimental replicates of independent wells of a differentiation; error bars represent SD. Statistical significance was determined using a one way-ANOVA with Tukey’s multiple comparisons test (panel C) and a Kruskal-Wallis test with Dunn’s multiple comparisons test (panel D, as data was non-parametric); *P < 0.05, ****P < 0.0001. **(E-G)** Transmission electron microscopy of M75 infection in iAECs at **(E)** 2000X and in iAT2s at **(F)** 2500X and **(G)** 1200X. Scale bar = 1 µm, scale bar in zoomed image = 200 nm.

### Virulence-independent *S. pyogenes* gene expression is dynamic across time and compartments in iPSC-derived lung epithelium

The upper respiratory tract represents a primary niche for *S. pyogenes* colonisation and local disease (pharyngitis) whereas invasive disease, including pneumonia, is an uncommon outcome. To explore whether bacterial gene expression changes in virulence and metabolic pathways underlie these divergent outcomes, we leveraged our iPSC-derived airway and alveolar models to recreate both niches and assessed global *S. pyogenes* gene expression across cell types and time (Figure 2A). Airway (iAECs) and alveolar (iAT2s) epithelial cells were infected with *S. pyogenes* (M1_UK_ strain) for 6 or 24 hours prior to RNA-seq analysis. Principal component analysis (PCA) revealed that infection of iAECs and iAT2s triggered divergent gene expression responses in *S. pyogenes*, particularly by 24 hours post infection (Figure 2B). Unbiased hierarchical clustering across all differentially expressed genes segregated samples by infection time point (Figure 2C). Consistent with this, differential gene expression (DEG) analysis of these time points revealed 244 and 283 DEGs between 6 and 24 hours in infected iAECs and iAT2s, respectively (Supplemental Figure 1A-D). We next directly compared *S. pyogenes* gene expression across airway or alveolar cell culture models at each time point. At 6 hours, 200 genes were differentially expressed between *S. pyogenes* infection of iAECs vs iAT2s (Figure 2D). By 24 hours, there were 338 differentially expressed genes distinguishing *S. pyogenes* grown in the iAT2 versus iAEC cultures (Figure 2E). Gene Ontology (GO) analysis showed statistically significant enrichment of terms associated with protein folding after 6 hours of infection by *S. pyogenes* in iAT2 cultures. At the 6 hour analysis point, no enriched GO terms reached a false discovery rate (FDR) <0.05 for iAECs (Figure 2F). In contrast, at 24 hours, no GO terms were significantly enriched after multiple testing correction (FDR < 0.05), suggesting that despite differential gene expression, *S. pyogenes* exhibited broadly similar functional pathways in airway and alveolar infections at 24 hours. *S. pyogenes* deploys a wide array of surface-associated and secreted virulence factors that shape interactions with host epithelial cells, immune defence and the transition to invasive disease (47). These include M protein (*emm*), superantigens (*speA*), proteases and immune-modulating enzymes (*speB, scpA, scpC/SpyCEP, sic*), tissue invasion (*ska*) and cytolytic toxins (*slo, sagA*). Expression of many virulence factors (*emm, slo, speA, sic, ska, scpA/B, sagA*) were highest after 6 hours of infection compared with 24 hours (Figure 2G). This was irrespective of the cell type being infected, with *S. pyogenes* in both iAEC or iAT2 cultures expressing similar levels of virulence genes. Finally, we explored whether *speB* expression varied depending on the *S. pyogenes* strain used to infect iPSC-derived lung epithelial cells and found that after 6 hours of infection, *speB* levels were comparable between the M1_UK_ or M75 strains irrespective of whether they infected iAECs or iAT2s (Figure 2H). Collectively, our results indicate that broad *S. pyogenes* transcriptional profiles change dynamically between 6 and 24 hours post infection in a cell type-dependent manner, without corresponding changes in virulence gene expression.

**Figure 2.**
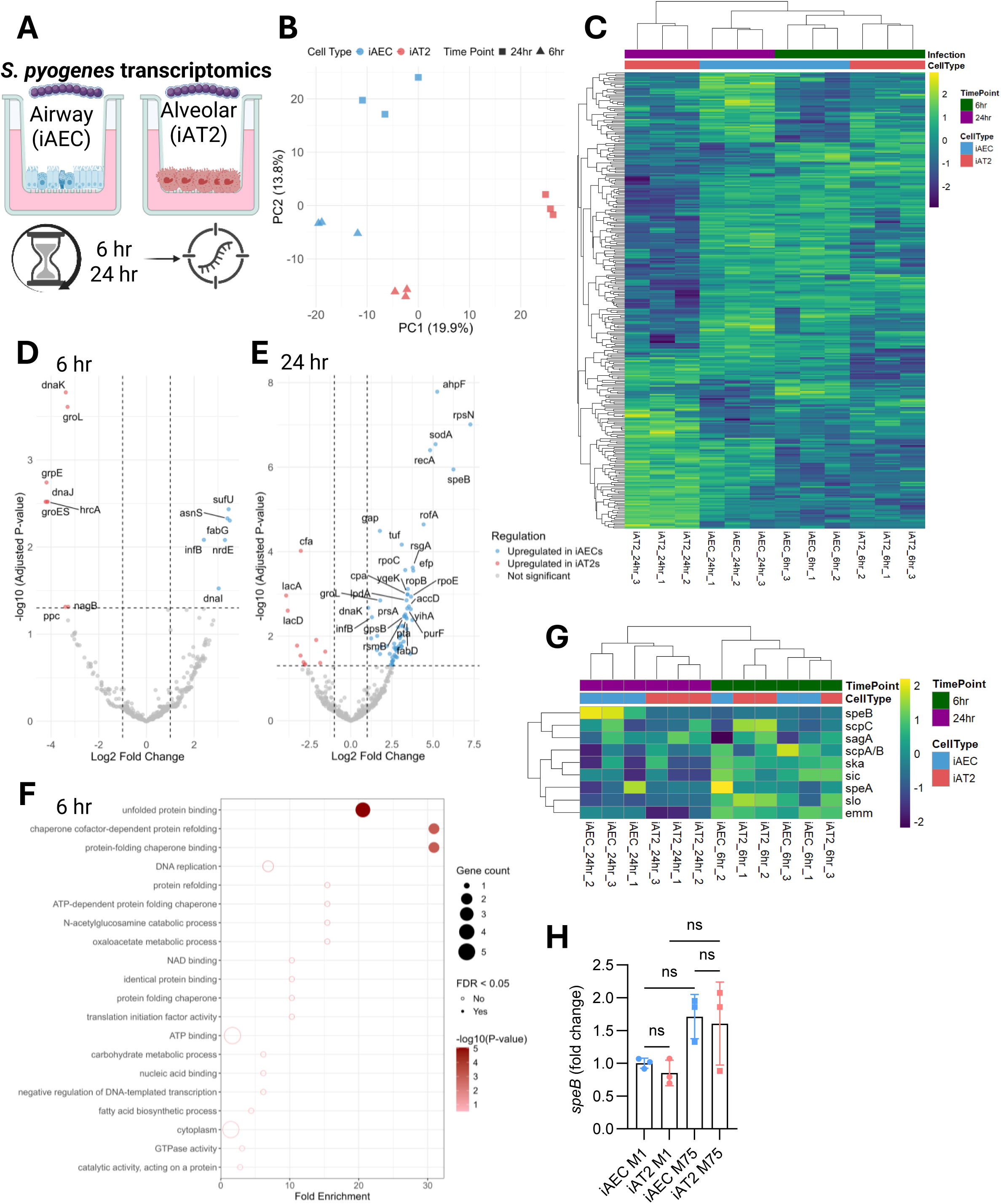
*S. pyogenes* gene expression is temporally dynamic and compartment specific in iPSC-derived lung epithelium. iAECs or iAT2s were infected with *S. pyogenes* M1_UK_ (1×10^6^ CFU/ml) for 6 or 24 hours. **(A)** Schematic overview of *S. pyogenes* transcriptomics. Human iPSCs underwent directed differentiation to airway or alveolar epithelial cells, which were further matured at air-liquid interface (ALI) prior to infection with *S. pyogenes*. Global *S. pyogenes* transcriptome was analysed by RNA-sequencing. **(B)** Principal component analysis (PCA) of bacterial gene expression at 6- or 24-hours post infection of iAECs or iAT2s. **(C)** Heatmap of differentially expressed genes (n=312). Expression values are row-scaled (z-score normalized) where colour represents relative expression from low (purple) to high (yellow). **(D)** Differentially expressed genes (DEGs) after 6 hours or **(E)** 24 hours infection of iAECs or iAT2s. **(F)** GO enrichment analysis was performed on differentially expressed genes (padj < 0.05, |log2FC| > 1) comparing iAT2 (red) vs iAEC at 6 hours. Enrichment was calculated using hypergeometric testing with Benjamini-Hochberg FDR correction. **(G)** Expression of virulence factors following 6 or 24 hours of iAEC or iAT2 infection, displayed by hierarchical clustering with row scaling (z-score normalised where colour represents relative expression from low (purple) to high (yellow)). **(H)** qRT-PCR expression of *speB* in iAECs or iAT2s infected with M1_UK_ or M75 strains for 6 hours. n = 3 experimental replicates of independent wells of a differentiation; error bars represent SD. Statistical significance was determined using a one way-ANOVA with Tukey’s multiple comparisons test.

### Immune responses diverge by lung epithelial cell type following *S. pyogenes* infection

We next sought to characterise the host immune response to *S. pyogenes* infection over time in airway (iAECs) and alveolar (iAT2) epithelial cells. In iAECs, infection triggered the up-regulation of genes encoding mediators of inflammation (*IL1B*) and chemokines (*CXCL8, CXCL2*) (Figure 3A and Supplementary Figure 2A-D). This response was largely replicated in cultures comprised of alveolar epithelial cells (Figure 3B). At the transcriptional level, the magnitude of host response was dependent on continued presence of bacteria, since host responses waned following removal of the inoculum (Supplementary Figure 2E-F). Moreover, this effect was observed upon infection with live, whole bacteria, whereas stimulation with lipopolysaccharide (LPS; Gram-negative component) or lipoteichoic acid (LTA; Gram-positive component) did not elicit a comparable immune response (Supplementary Figure 2G-L). We next assessed infection of the models with different strains of *S. pyogenes*. At 6 hours, we found infection with the M1_UK_ strain induced a stronger proinflammatory response (*IL6, IL1B, TNF*) than the M75 strain in both cell types (Figure 3C). Furthermore, some mediators were not upregulated by infection of iAT2s (e.g., *CXCL8, CCL2*) at 6 hours. We next assessed cytokine and chemokine secretion basolaterally by each cell type following 6 hours of infection with either the M1_UK_ or M75 strain. Of the 17 proteins detected in the assay, iAECs secreted significantly higher levels of cytokines and chemokines than iAT2s, both at baseline (uninfected) and following infection with either strain (Figure 3D). Infection significantly increased iAEC secretion of IL-6 in response to M1_UK_ and G-CSF in response to M75 (Figure 3D and Supplementary Figure 2M–N). In contrast, iAT2s exhibited minimal cytokine and chemokine secretion under both basal and infected conditions, with only VEGF significantly increased following M1_UK_ infection (Figure 3D and Supplementary Figure 2O). Given the limited inflammatory response of iAT2s, we next examined whether differential cellular responses to infection were instead reflected in cell viability and death by determining cells in early and late apoptosis using flow cytometry. Following infection with M1_UK_, a significantly greater proportion of iAT2s were in late apoptosis at both 6 and 24 hours compared with uninfected controls (Figure 3E and Supplementary Figure 2P). In contrast, no significant differences were observed in the proportions of viable, early apoptotic, or late apoptotic iAECs at either time point relative to uninfected controls (Figure 3E and Supplementary Figure 2P). Together, these data demonstrate distinct epithelial cell-type specific responses to *S. pyogenes* infection, with iAECs mounting a robust inflammatory response, while iAT2s exhibit limited cytokine/chemokine induction but increased susceptibility to cell death. Lastly, because our alveolar cultures lacked type 1 cells, we generated iPSC-derived type 1 alveolar epithelial cells at ALI (48) and infected them with the M1_UK_ strain. Compared with the type 2 alveolar epithelial cultures, these cells exhibited an even weaker host response at 6 and 24 hours post-infection, with only *CXCL8* induction observed (Supplementary Figure 2Q-T).

**Figure 3.**
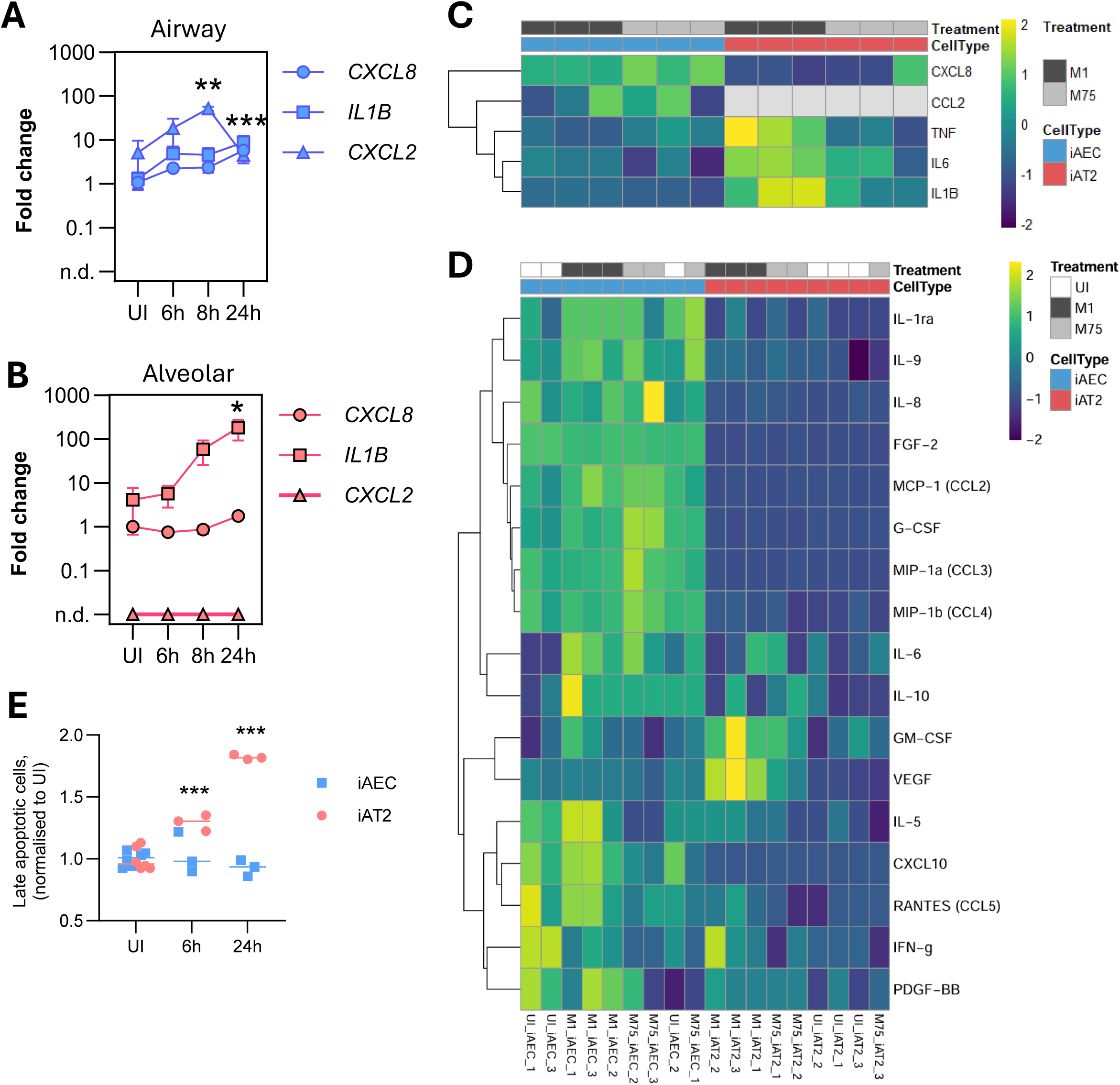
Host response to *S. pyogenes* infection is compartment and strain specific. **(A)** iAECs or **(B)** iAT2s were infected with *S. pyogenes* M1_UK_ (1×10^6^ CFU/ml) for 6, 8 or 24 hours. Gene expression was assessed by qRT-PCR. n.d. = no detected. **(C)** iAECs/iAT2s were infected with *S. pyogenes* M1_UK_ or M75 (1×10^6^ CFU/ml) for 6 hours. Gene expression was assessed by qRT-PCR. Heatmap representation with row scaling (z-score normalised where colour represents relative expression from low (purple) to high (yellow). Grey indicates not detected). **(D)** iAECs/iAT2s were infected with *S. pyogenes* M1_UK_ or M75 (1×10^6^ CFU/ml) for 6 hours. Basolateral media was analysed using a multiplexed cytokine assay. Heatmap representation of cytokine levels displayed by hierarchical clustering with row scaling (z-score normalised where colour represents relative expression from low (purple) to high (yellow)). **(E)** iAECs/iAT2s were infected with *S. pyogenes* M1_UK_ (1×10^6^ CFU/ml) for 6 or 24 hours. Late apoptotic cells (Annexin V+ 7-AAD+) were analysed by flow cytometry and normalized to uninfected (UI) cells. n = 3 experimental replicates of independent wells of a differentiation; error bars represent SD. Statistical significance was determined using a two way-ANOVA with Tukey’s multiple comparisons test; *P < 0.05, **P < 0.01, ***P < 0.001.

### Airway epithelial cells promote inflammatory signalling following *S. pyogenes* infection

Given the response of human airway epithelium to *S. pyogenes* infection has not been explored, we next assessed global transcriptomic responses in our iPSC-derived airway epithelial cells grown at ALI after 6 and 24 hours of infection with *S. pyogenes* (M1_UK_) (Figure 4A). PCA found that at each infection time point and the control (uninfected iAECs) clustered separately (Figure 4B). Gene expression analysis revealed 79 and 203 differentially expressed genes (DEGs) relative to uninfected control iAECs at 6 and 24 hours, respectively (Figure 4C). There was significant overlap in DEGs at these time points (Figure 4D). Several leukocyte-attracting chemokines, including *CXCL1* and *CCL20*, were amongst the top DEGs (Figure 4E). Gene set enrichment analysis (GSEA) and gene regulatory network (GRN) analysis demonstrated an enrichment in hypoxia and interferon responses, with NFKB2 and REL identified as central transcriptional regulators with high connectivity in the gene regulatory network, particularly at 24 hours post infection (Figure 4F-H and Supplemental Figure 3A-B). During infections, adaptive and innate immune cells preferentially use glycolysis during infections to supply the required energy for effective immune responses (49). In our dataset we observed a shift in metabolic activity of the airway epithelium; while uninfected iAECs predominantly utilise oxidative phosphorylation, *S. pyogenes* infected cells upregulated genes encoding key components of the glycolysis pathway (Figure 4F-H). Finally, we assessed the levels of 27 secreted cytokines and chemokines in the basolateral compartment of iAECs after 6 and 24 hours of infection with *S. pyogenes* (M1_UK_). IL-1RA was the only protein that increased significantly over the course of infection, whereas RANTES, IL-8, and CXCL10 levels trended lower by 24 hours post-infection (Figure 4I and Supplemental Figure 3C-D). Together, our transcriptome and secretome profiling indicates that *S. pyogenes* infection of the human pseudostratified airway epithelium triggers rapid and dynamic innate immune responses coordinated by NF-κB-signalling.

**Figure 4.**
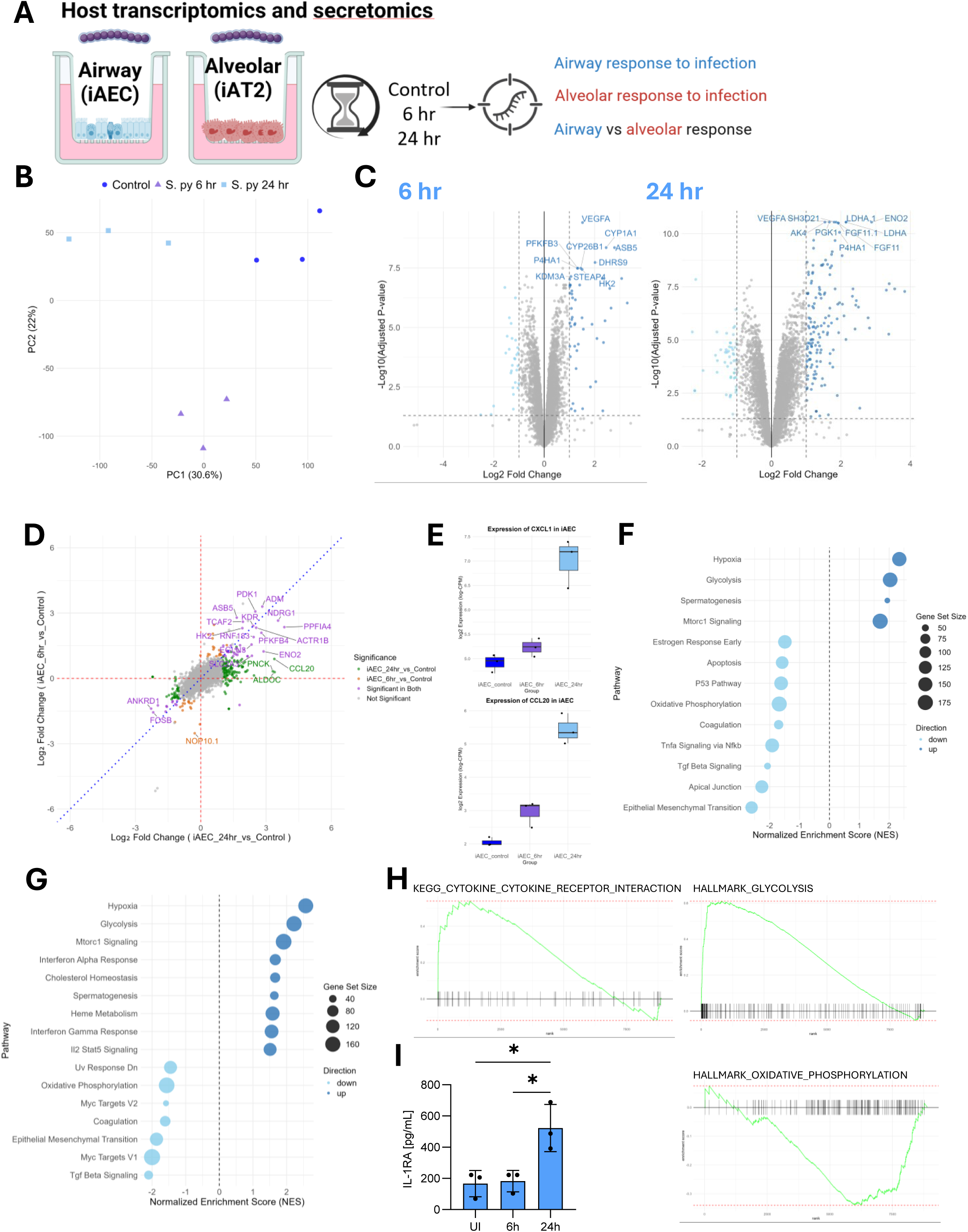
iPSC-derived airway epithelial cells induce inflammatory signalling in response to *S. pyogenes* infection. iPSC-derived AECs were infected with *S. pyogenes* M1_UK_ (1×10^6^ CFU/ml) for 6 or 24 hours. **(A)** Schematic overview of host transcriptomics. Human iPSCs underwent directed differentiation to airway or alveolar epithelial cells, which were further matured at air-liquid interface (ALI) prior to infection with *S. pyogenes*. Global host transcriptomes were analysed by RNA-sequencing **(B)** Principal component analysis (PCA) of uninfected (control), 6-hour or 24-hour infection iAECs. **(C-D)** Differentially expressed genes (DEGs) after 6 hours or 24 hours of *S. pyogenes* infection, compared with uninfected controls. **(E)** Expression of chemokines. **(F)** Gene set enrichment analysis (GSEA) using the Hallmark database of 6 hours *S. pyogenes* infection or **(G)** 24 hours *S. pyogenes* infection compared with uninfected controls. **(H)** Enrichment plots of differentially expressed pathways after 24 hours of *S. pyogenes* infection using the KEGG and Hallmark databases, relative to uninfected controls. **(I)** IL-1RA levels were measured using a multiplexed cytokine assay. n = 3 experimental replicates of independent wells of a differentiation; error bars represent SD. Statistical significance was determined using a one way-ANOVA with Tukey’s multiple comparisons test; *P < 0.05, **P < 0.01, ***P < 0.001.

### Proinflammatory and proliferative responses dominate type 2 alveolar epithelial responses to *S. pyogenes* infection

Since AT2s are central to pneumonia pathogenesis (50), we next examined transcriptomic changes triggered by *S. pyogenes* (M1_UK_) infection in iPSC-derived AT2s grown at ALI. iAT2s exposed to *S. pyogenes* for 6 or 24 hours and control (uninfected) conditions clustered distinctly on a PCA plot (Figure 5A). Infection triggered a rapid transcriptomic response (702 DEGs), which appeared to abate by 24 hours (49 DEGs) (Figure 5B). In contrast with airway epithelial infection, there were fewer concordant DEGs between the two infection time points in iAT2s (Figure 5C). GSEA found that *S. pyogenes* infection upregulated proinflammatory (e.g., TNF-α, IL-6 JAK/STAT) and proliferation (e.g., E2F, Myc, G2M checkpoint) pathways in iAT2s (compared with uninfected controls) at both 6 and 24 hours (Figure 5D). Indeed, we discovered significant enrichment in E2F targets (Figure 5E and Supplemental Figure 4A). Furthermore, expression of key genes in the cell cycle changed dynamically over infection; *PCNA* (S phase) peaked at 6 hours, *TOP2A* (G2/M) peaked at 24 hours and *MKI67* (G1-M) steadily increased with time (Figure 5F), coinciding with the temporal changes in apoptosis observed in iAT2s (Figure 3). TNF-α signalling via NF-κB was also significantly enriched (Figure 5G and Supplemental Figure 4A-B), including downstream targets (e.g., *DUSP5, CXCL8*) (Figure 5H) following *S. pyogenes* infection of iAT2s. Interestingly, despite this transcriptional signature of inflammation, iAT2s exhibited minimal cytokine and chemokine secretion at both 6 and 24 hours following M1_UK_ infection. Instead, levels of several chemokines, including IL-8 and CXCL10, were significantly reduced (Figure 5I and Supplementary Figure 4C–D). Taken together, these data indicate that *S. pyogenes* infection of the alveolar epithelium induces robust proinflammatory and proliferative gene expression programs without a corresponding increase in cytokine and chemokine secretion in this infection window.

**Figure 5.**
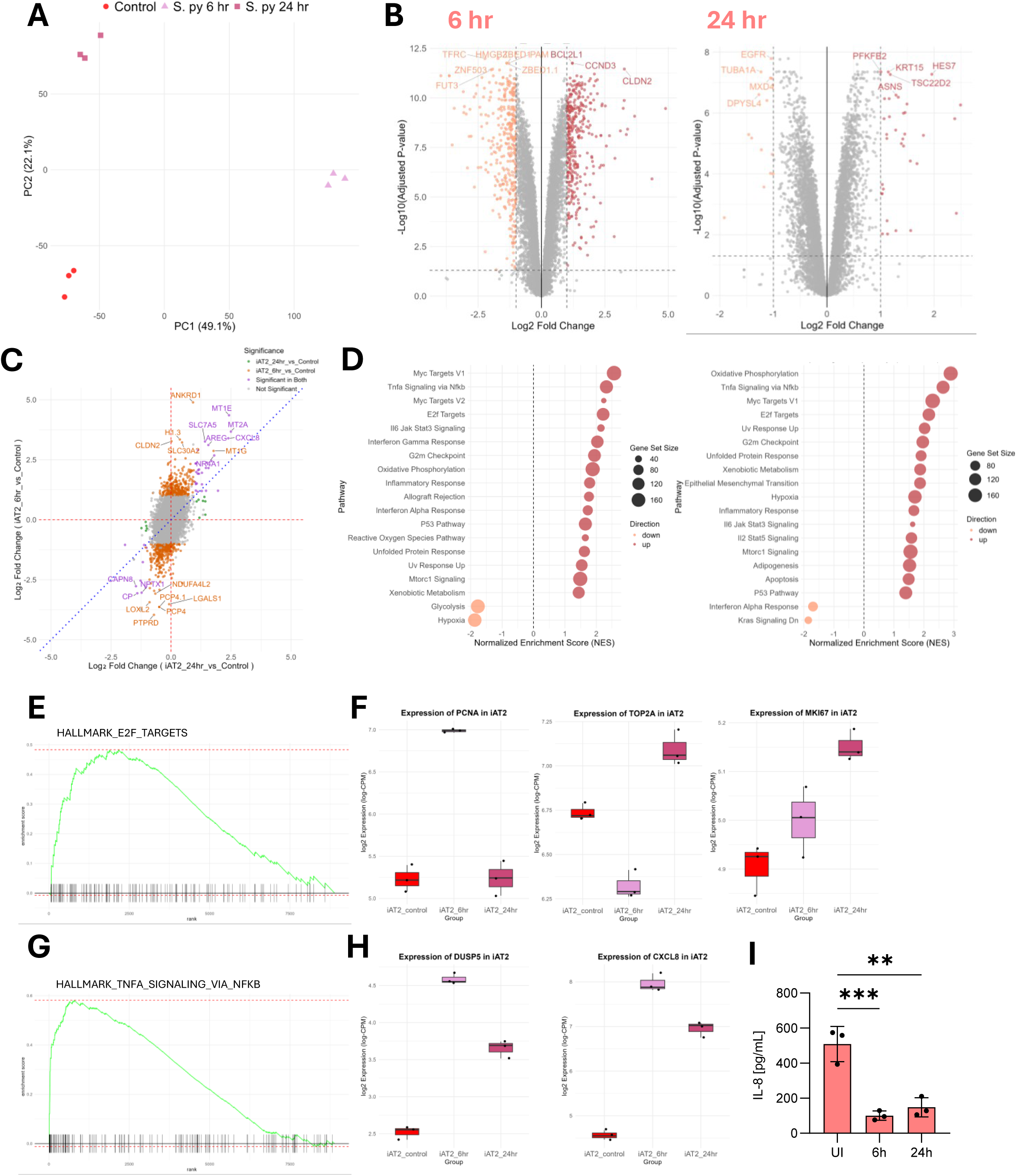
iPSC-derived alveolar epithelial cells show proinflammatory and proliferative signalling following *S. pyogenes* infection. iPSC-derived AT2s were infected with *S. pyogenes* M1_UK_ (1×10^6^ CFU/ml) for 6 or 24 hours. **(A)** Principal component analysis (PCA) of uninfected (control), 6-hour or 24-hour infection iAT2s. **(B-C)** Differentially expressed genes (DEGs) after 6 hours or 24 hours of *S. pyogenes* infection, compared with uninfected controls. **(D)** Gene set enrichment analysis (GSEA) using the Hallmark database of 6 hours *S. pyogenes* infection (left) or 24 hours *S. pyogenes* infection (right) compared with uninfected controls. **(E)** Enrichment plots of Hallmark E2F targets, upregulated in iAT2s at 24 hours of *S. pyogenes* infection. **(F)** Expression of proliferation-associated genes. **(G)** Enrichment plots of Hallmark TNF-a signalling via NFkB, upregulated in iAT2s at 24 hours of *S. pyogenes* infection. **(H)** Expression of proinflammatory genes. **(I)** IL-8 levels were measured using a multiplexed cytokine assay. n = 3 experimental replicates of independent wells of a differentiation; error bars represent SD. Statistical significance was determined using a one way-ANOVA with Tukey’s multiple comparisons test; *P < 0.05, **P < 0.01, ***P < 0.001.

### Limited interferon signalling in type 2 alveolar cells permits *S. pyogenes* invasion and is rescued by IFN-β

Given the range of clinical outcomes of *S. pyogenes* infection in the respiratory tract, from asymptomatic colonisation to pneumonia, we hypothesised that regional differences in host epithelial responses could contribute to this variability. Direct comparison of *S. pyogenes*-infected alveolar (iAT2s) with airway (iAECs) epithelial cells, after normalisation for cell type, revealed 1062 DEGs at 6 hours and 382 DEGs at 24 hours post infection (Figure 6A and Supplemental Figure 5A-C). Notably, relatively few DEGs overlapped between the two cell types at either time point (Figure 6B). GSEA revealed significant divergence in pathways. At 24 hours post *S. pyogenes* infection, iAECs were significantly enriched for hypoxia, glycolysis and IFN-α response (Figure 6C-D). In contrast, enriched pathways in iAT2s included oxidative phosphorylation, epithelial-to-mesenchymal transition and TNF-α signalling via NF-κB (Figure 6C-D).

**Figure 6.**
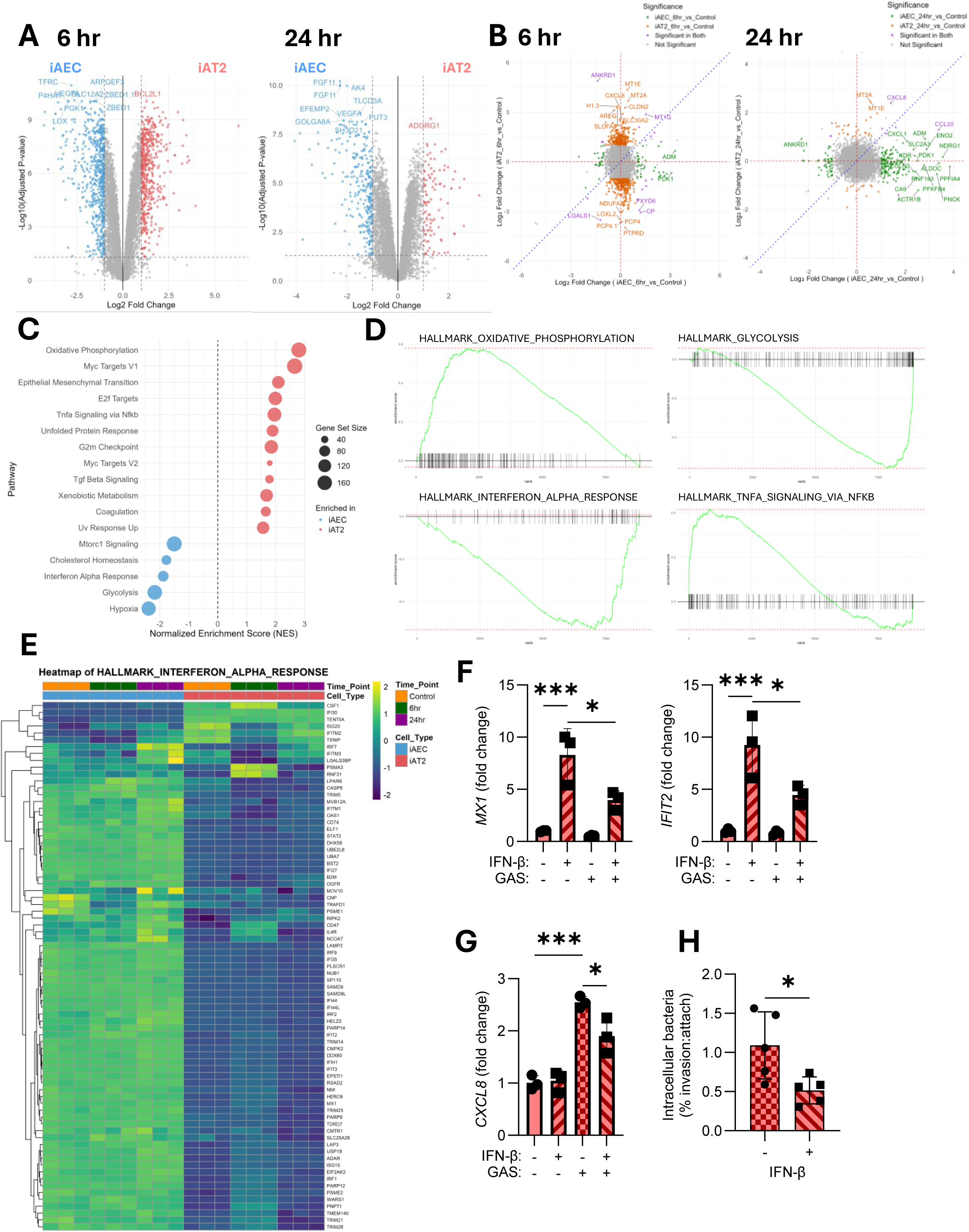
iPSC-derived alveolar epithelial cells show limited baseline interferon signalling which is overcome by IFN-β to restrict *S. pyogenes* invasion. iPSC-derived AECs and AT2s were infected with *S. pyogenes* M1_UK_ (1×10^6^ CFU/ml) for 6 or 24 hours, bulk RNA-seq analysis performed and then gene expression changes were normalized to uninfected controls to control for cell type. **(A-B)** Differentially expressed genes (DEGs) upregulated in iAT2s after 6 hours or 24 hours of *S. pyogenes* infection compared to iAECs. **(C)** Gene set enrichment analysis (GSEA) using the Hallmark database after 24 hours *S. pyogenes* infection in iAT2s (red) versus iAECs (blue). **(D)** Enrichment plots of Hallmark pathways upregulated in iAT2s compared to iAECs after 24 hours of *S. pyogenes* infection. **(E)** Expression of genes in Hallmark IFN-α response gene set. Expression values are row-scaled (z-score normalized) where colour represents relative expression from low (purple) to high (yellow). **(F)** iAT2s were treated with IFN-β and infected with *S. pyogenes* (M1_UK_) for 24 hours. Expression of IFN-stimulated genes (*MX1, IFIT2*) and **(G)** proinflammatory (*CXCL8*) genes were measured via qRT-PCR. **(H)** iAT2s were treated with IFN-β 24 hours prior to infection with *S. pyogenes* (M1_UK_) for 6 hours. Bacterial attachment and invasion were quantified via CFU assay. n = 3 experimental replicates of independent wells of a differentiation; error bars represent SD. Statistical significance was determined using a one way-ANOVA with Tukey’s multiple comparisons test or unpaired t-test (panel H); *P < 0.05, ***P < 0.001.

We were intrigued that an IFN-α response was consistently associated with infection responses for iAECs but not iAT2s. Despite normalising for cell type in our analyses, many genes associated with IFN signalling were expressed at baseline in iAECs, but not iAT2s (Figure 6E). Moreover, expression of the type I IFN receptor (*IFNAR1/2*) was highly expressed in iAECs, but not iAT2s, while the type III IFN receptor (*IFNLR1* and *IL10RB*) is similarly expressed on both cell types (Supplemental Figure 5D). These data suggest that iAECs but not iAT2s are primed and responsive to type I IFN. Studies in mice suggest that type I IFN protects against invasive *S. pyogenes* infection in other organs (51, 52), and *S. pyogenes* can restrict type I IFN signalling (53). Since these studies suggest that a type I IFN response is beneficial in host immune responses to *S. pyogenes,* we next determined whether type I IFN supplementation could protect iAT2s during infection. IFN-β was added prior to *S. pyogenes* infection (M1_UK_) of iAT2s. Exogenous IFN-β had no effect on *S. pyogenes* attachment and expression of the virulence gene, *speB,* nor did it protect iAT2s from losing barrier integrity following infection (Supplemental Figure 5E-G). The expression of IFN-stimulated genes (ISGs) (*MX1* and *IFIT2*) were significantly elevated by IFN-β treatment, although this induction was blunted when given in conjunction with infection, suggesting that *S. pyogenes* infection interferes with IFN signalling in iAT2s (Figure 6F). Treatment with IFN-β significantly reduced *CXCL8* induction following *S. pyogenes* infection of iAT2s (Figure 6G), consistent with IFN-β treatment limiting bacterial invasion of iAT2s (Figure 6H).

## DISCUSSION

Here, we have established a novel, scalable human model for *S. pyogenes* respiratory infection. Our findings highlight how epithelial context shapes the outcome of host–pathogen interactions. We used iPSC-derived airway and alveolar epithelial models to define how distinct lung compartments respond to infection with M1_UK_ and M75 strains. Both strains adhered to and invaded epithelial cells, with differences linked to strain and epithelial type. Transcriptomic profiling revealed compartment-specific host responses, with type I IFN responses more robustly primed in the airway, while IFN-supplementation restricts *S. pyogenes* invasion and inflammation in alveolar epithelial cells.

Study of the pathogenesis of *S. pyogenes* respiratory disease, particularly pneumonia, is difficult as the bacteria is a strict human pathogen and because human alveolar epithelial cells are difficult to obtain and are prone to *ex vivo* senescence (54). Moreover, sourcing airway and alveolar epithelial cells from the same donor is extremely challenging, introducing donor variability as a confounding factor when trying to compare the responses of airway and alveolar cells to specific challenges. To overcome these limitations, we used well-established ALI platforms representing the human airway and alveolar epithelium derived from iPSCs (28, 31, 40). These have been extensively benchmarked against *in vivo* counterparts, recapitulating key airway and alveolar functions (28–30, 36, 40), with their fidelity underscored by SARS-CoV-2 studies, where immune responses observed in iAT2s (32, 36) were subsequently reproduced *ex vivo* and *in vivo* (36, 54). Herein our findings establish iPSC-derived airway and alveolar epithelial models as a novel, physiologically relevant system for studying respiratory *S. pyogenes* infection.

Although *S. pyogenes* is primarily an extracellular pathogen, it can invade and replicate inside host cells (55). One proposed mechanism for severe *S. pyogenes* infections is invasion of the alveolar epithelium, enabling spread into the pleural cavity or bloodstream (56). Although uncommon, we observed consistently more *S. pyogenes* invasion of iPSC-derived alveolar epithelial cells, compared with the airway epithelium. Moreover, following invasion of alveolar epithelial cells we observed hyaluronic acid capsule shedding by *S. pyogenes*. Poorly encapsulated or capsule-mutant *S. pyogenes* invade immortalized skin and pharyngeal cells more efficiently (57, 58), suggesting that while the capsule protects against phagocytosis, it may impede entry into non-phagocytic cells. Certain *S. pyogenes* strains are strongly associated with more invasive disease, acquiring and expressing regulators of virulence, and superantigens, a family of virulence factors that stimulate expression of pro-inflammatory cytokines (17–19, 59). However, the full spectrum of *S. pyogenes* virulence gene expression in the lower respiratory tract remains unknown, as sampling the lungs during infection is rare. Our transcriptomic analysis comparing *S. pyogenes* following infection of iAECs or iAT2s revealed similar virulence gene expression early in infection, at a time point when invasion of iAT2s had already occurred. Collectively, our findings indicate that expression of virulence genes is not the main driver of susceptibility by iAT2s to infection.

Epithelial immune responses play a central role in restricting respiratory bacterial infection, both through intrinsic antimicrobial activity and by orchestrating the recruitment of immune cells. In contrast, *S. pyogenes* can provoke excessive inflammation via potent streptococcal pyrogenic exotoxins and virulence factors such as SpeA and SpeB, suggesting that the balance between effective defence and immune overactivation is critical to disease outcome. Our transcriptomic and secretomic analysis of iAECs and iAT2s following *S. pyogenes* infection revealed dynamic inflammatory signalling, particularly induction of chemokines, cytokine release and transcription factor networks dominated by regulators of inflammation (e.g., NF-κB/REL). Although there are no published *in vivo* datasets of lower respiratory *S. pyogenes* infections, our data is consistent with hyperinflammatory responses observed in human challenge samples (60), in primate bacteraemia studies (61), and *in vitro* studies using primary (e.g., tonsil epithelial cells) and immortalised lung epithelial cell lines (e.g., A549, HEp-2) (62–64).

Type I IFNs are potent inducers of antiviral immunity but can have paradoxical and context-dependent effects during bacterial infections. In mice, type I IFN signalling protects against invasive *S. pyogenes* infection of soft tissue (51, 52), and *S. pyogenes* can restrict type I IFN signalling by impairing plasmacytoid dendritic cell recruitment (53). These studies suggest a type I IFN response is beneficial in host immune responses to *S. pyogenes*, through modulation of myeloid cell function (65), including augmented *S. pyogenes* killing (66), and dampened inflammation. Unlike myeloid cells, our findings show that *S. pyogenes* did not induce IFN expression in lung epithelial cells. However, at baseline, iPSC-derived airway epithelial cells exhibit significantly higher protective type I IFN signalling than iAT2s. Consistent with this, *S. pyogenes* invasion of the alveolar epithelium was reduced only when type I IFN was provided exogenously. Together, these findings suggest that elevated basal type I IFN activity in airway epithelial cells may limit *S. pyogenes* invasion, complementing other airway defences such as mucous production and ciliary clearance. Moreover, our findings suggest that increased type I IFN signalling, either directly or through limiting *S. pyogenes* invasion, attenuates pro-inflammatory gene induction. For instance, *CXCL8* expression was restricted in IFN-β-treated iAT2s and was induced to a lesser extent following infection with the less invasive M75 strain compared with M1_UK_. Finally, while ISGs were upregulated when IFN-β was added to infected cultures, this response was weaker than in uninfected controls. Moreover, secretion of the ISG, CXCL10, was consistently reduced in both *S. pyogenes*-infected iAECs and iAT2s. Collectively, this suggests that *S. pyogenes* may actively suppress IFN-stimulated signalling in the lung epithelium. Similar suppression of IFN signalling has been reported in macrophages, where *S. pyogenes*-induced IL-10 limits IFN responses (67). Alternatively, enhanced invasion and associated cellular stress/death in iAT2s may impair appropriate IFN signalling. Future studies should define the mechanisms by which *S. pyogenes* restricts IFN signalling in the alveolar epithelium, with potential implications for host-directed therapies for pneumonia.

Our study is limited by the inclusion of only two strains of *S. pyogenes*. These strains were selected to reflect divergent bacterial and clinical phenotypes: M1 is considered a “throat-specialist” strain, is the most prevalent globally across all infections, and is disproportionately represented in invasive disease (17), while M75 is regarded as a more “generalist” strain, occurring in both throat and skin infections and is less frequently associated with severe/invasive disease than M1 (22, 68). However, we acknowledge that our study could not capture the full diversity of *S. pyogenes* nor the complexity of the relationships between strain variation and clinical phenotypes (15). M1_UK_ and M75 strains have been used in recent and ongoing controlled human challenge studies (22, 69), and these will provide valuable future references for comparing our findings using iPSC-derived models. Moreover, our analysis of host-pathogen transcriptomic responses was performed in bulk, which cannot resolve the heterogeneity between host cells directly infected with *S. pyogenes* and neighbouring uninfected cells. The ability to capture both host and bacterial transcriptional responses at single-cell resolution, as recently demonstrated for *Bacillus subtilis* (70), would represent a major advance for the *S. pyogenes* field. Finally, future work could extend these models by incorporating innate immune cells, as we have recently done with macrophages (37, 38).

In sum, using iPSC-derived airway and alveolar models, we demonstrated that host responses vary by strain and lung cell type. Type I IFN is more strongly induced in the airway epithelium, and exogenous IFN-β protects the alveolar epithelium from *S. pyogenes* invasion. Together, these results define a scalable human model and demonstrate that epithelial context critically shapes *S. pyogenes* infection dynamics.

## MATERIALS AND METHODs

### Human iPSC maintenance

iPSCs used in this study consist of the BU3 NGST CRISPRi (71) and SCT3010 (38). iPSCs required daily media change with StemFlex, mTeSR or mTeSR+ media. Ethical approval for the generation and/or use of human iPSCs were obtained from Murdoch Children’s Research Institute (MCRI) and Boston University and were carried out in accordance with the National Health and Medical Research Council of Australia (NHMRC) regulations.

### Directed differentiation of iAECs and iAT2s

Human iPSC lines were directed to differentiate into either iAECs or iAT2s, as previously described (28, 72). In brief, iPSCs were differentiated to definitive endoderm using the STEMdiff Definitive Endoderm Kit (StemCell Technologies, 05210), and flow cytometry was used to verify CXCR4+ cKit+ expression. Cells were then dissociated using Gentle Cell Dissociation Reagent (StemCell Technologies, 100-0485) and replated on plates coated with growth factor-reduced Matrigel (Corning, 354277). Afterwards, cells were cultured in the anteriorisation media “DS/SB” (cSFDM with 2 µM Dorsomorphin (Tocris, 3093/10) and 10 µM SB431542 (Tocris, 1614)) for 3 days, with the first 24 hours also supplemented with 10 µM Y-27632 (Tocris, 1254). From day 6 of the differentiation, Cells were kept in “CBRa” (cSFDM with 3 µM CHIR99021 (Tocris, 4423), 10 ng/ml rhBMP4 (R&D Systems, 314-BP), and 100 nM Retinoic Acid (Sigma, R2625)) to induce NKX2-1^+^ lung progenitors. On days 14 or 15, cells were sorted for NKX2-1^+^ lung progenitors using a FACSAria Fusion (BD Biosciences). Sorted NKX2-1^+^ lung progenitors were embedded in growth factor-reduced Matrigel (Corning, 356230) droplets and maintained in either “2,10+DCIY” (cSFDM with 250 ng/ml FGF2 (R&D Systems, RDS233FB500), 100 ng/ml FGF10 (PeproTech, AF-100-26), 50 nM Dexamethasone (Sigma, D4902), 0.1 mM 8BrcAMP (Sigma, B7880), 0.1 mM IBMX (Sigma, I5879), and 10 µM Y-27632) or “CK+DCI” (cSFDM with 3 µM CHIR99021, 10 ng/ml rhKGF (R&D Systems, RDS251KG050), 50 nM Dexamethasone, 0.1 mM 8BrcAMP, and 0.1 mM IBMX). 2,10+DCIY or CK+DCI media was replaced every 2-3 days.

On day 30, iBCs were passaged, embedded in Matrigel droplets once more, and supplemented with 2,10+DCIY. The following day, 2,10+DCIY was replaced with Basal Cell Media (PneumaCult ExPlus (Stem Cell Tech, 05040) with 1 µM A83-01 (Tocris, 2939), 1 µM DMH1 (Tocris, 41-261-0), 10 µM Y-27632, and 200 nM Hydrocortisone (Stem Cell Technologies, 74142), which was replaced every 2-3 days. iBCs were serially passaged every 2 weeks and re-sorted when needed based on NKX2-1^GFP^+ NGFR+ expression or NGFR+ EpCAM+. After each passage, cells were supplemented with Basal Cell Media.

iAT2s were serially passaged every 2 weeks and re-sorted when needed based on NKX2-1^GFP^+ SFTPC^tdTomato^+ expression or CPM+. After each passage, cells were supplemented with CK+DCI+Y (CK+DCI with Y-27632).

### Air-liquid interface culture of iAECs and iAT2s

100 µl 2D Matrigel (diluted in DMEM/F12) was added to the apical surface of each 6.5 mm Transwell (Stem Cell Technologies, 38024). Transwells were incubated at 37°C for 30 minutes to let the Matrigel set, then excess 2D matrigel was aspirated. 100 µl DMEM was added to wash the apical surface and aspirated. Immediately after, 2000 cells/µl of iBCs or iAT2s, resuspended in 100 µl media (Basal Cell Media or CK+DCI+Y), were plated into the apical chamber. 500 µl of the same media was added to each of the basolateral chambers. For iBCs, once confluency was confirmed, both apical and basolateral media were replaced with ALI maintenance media (PneumaCult ALI Complete Basal Medium (Stem Cell Tech, 05002) with 100X PneumaCult-ALI Maintenance supplement (Stem Cell Tech, 05001), 4 µg/ml Heparin solution (Stem Cell Tech, 07980), and 1 µM Hydrocortisone). The next day, apical media was aspirated. For iAT2s, 48 hours post-plating, apical media was aspirated and basolateral CK+DCI+Y was replaced with CK+DCI. Following the formation of an air-liquid interface, basolateral media was replaced every 2-3 days. 3-4 weeks post-airlift, the airway iBCs at ALI would have differentiated and matured into iAECs (consisting of multiciliated, club, goblet, and basal epithelial cells in a pseudostratified epithelium) and would be ready for experiments (28). iAT2 ALIs were ready for experiments 5-7 days post-airlift (73).

To create iPSC-derived type 1 alveolar epithelial cells (iAT1s), 200,000 iAT2s were plated on 6.5 mm Transwells. Media was immediately switched to cSFDM containing 10 mM LATS-IN-1 (MedChemExpress, HY-138489), 50 nM Dexamethasone, 0.1 mM 8BrcAMP, and 0.1 mM IBMX (“L-DCI” media), as described (48).

Transepithelial electrical resistance in ALI cultures was measured using a Millicell ERS-2 Voltohmmeter (Merck, MERS00002).

### *Streptococcus pyogenes* culture and infection

*S. pyogenes* strain M1_UK_ (Strain ID MCRI0102) was isolated from a patient in the ASAVI STAMPS study, and strain M75 (Strain ID 611024) (74, 75) was isolated from the throat of a patient with paediatric pharyngitis in the Royal Children’s Hospital, Melbourne. Bacteria were streaked onto horse blood agar (HBA), incubated overnight at 37°C in 5% CO₂, and the next day, single colonies were used to inoculate Todd Hewitt broth + 1% (w/v) yeast extract (THY). Mid-logarithmic cultures were pelleted, resuspended in THY + 10% (v/v) glycerol, and stored at −80°C.

Frozen stocks were thawed at room temperature for 5 min, centrifuged at 5,000 × *g* for 5 min, and resuspended in CK+DCI (no primocin) or ALI maintenance media (no primocin) to a concentration of 1×10^6^ colony forming units per ml (CFU/ml). 100µl of bacteria (1×10^5^ CFU) or media-only controls (CK+DCI or ALI maintenance media; no primocin) were added to each apical chamber of iAEC or iAT2 ALIs. Plates were centrifuged at 200 × *g* for 5 min, and incubated at 37°C in 5% CO₂. Lipopolysaccharide (LPS; 1 µg/ml) or lipoteichoic acid (LTA; 0.1–1 µg/ml) were used as Gram-negative or Gram-positive mimetics, respectively, and were likewise added apically into 100 µl media for the desired duration.

Following infection, bacteria were removed, and apical chambers were washed with phosphate-buffered saline (PBS). Apical chambers of “invasion” samples were treated with 200 µg/ml Gentamicin (ThermoFisher, 15710-064) for 1 h. Basolateral media were snap-frozen and stored. After gentamicin (for “invasion”) and after washing with PBS, cells were dissociated with Accutase (Stem Cell Technologies, 7922) and lysed with 0.025% (v/v) Triton-X100 (Sigma-Aldrich, 9036-19-5). Bacteria were enumerated from “total attachment” (no gentamicin) or “invasion” groups by serial dilution and drip plating of 5 µl onto HBA (Horse Blood agar) or 10 µl onto TTC-THY (triphenyltetrazolium chloride with Todd-Hewitt Broth) agar. Plates were incubated overnight at 37°C in 5% CO₂. Mean CFUs from acceptable dilutions (5-50 CFU/track) were adjusted for dilution and expressed as CFU/ml. Percentage (%) of bacterial inoculum was calculated from “total attachment” relative to the initial inoculum, and % invasion determined using “invasion” relative to “total attachment.”

### Flow cytometry and cell sorting

On day 3 of the differentiation, endoderm cells were stained for CXCR4 (BD Bioscience, 555974) and c-Kit (Biolegend, 313206) and analysed using a Fortessa flow cytometer (BD Biosciences). NKX2-1+ cells were isolated based on NKX2-1^GFP^ expression, or CD47 and CD26 expression. For the latter, cells were stained with antibodies (CD47^PerCPCy5.5^ (Biolegend, 323110) and CD26^PE^ (Biolegend, 323110)) for 30 minutes on ice, and CD47^hi^CD26^lo^ cells were isolated by FACS (27). Where indicated, NGFR+ iBCs were purified by FACS on the basis of an NGFR^APC^+ (Biolegend, 345105) stain or based on EpCAM (Biolegend, 324206) expression. Whereas SFTPC+ iAT2s were purified by FACS on the basis of a SFTPC^tdTomato^ reporter gene (72) or on the basis of CPM expression (Novachem, 014-27501) (73). Cells were resuspended in FACS buffer: Hank’s Balanced Salt Solution (ThermoFisher) with 2% FCS in PBS and 10 µM Y-27632. Live cells were sorted using calcein blue, Zombie NIR or Zombie R718 (Table 4). Cells were isolated using a FACS Aria (BD Biosciences) at the Murdoch Children’s Research Institute Flow Cytometry Core Facility.

To identify viable and apoptotic cells, samples were stained with antibodies against Annexin V (Biolegend, 640924 or BD, 563972) and 7-AAD (Biolegend, 640934) in Annexin V staining buffer (Biolegend, 640934) and analysed using a LSRFortessa X-20 flow cytometer (BD Biosciences). 7-AAD- Annexin V- cells were viable, 7-AAD- Annexin V+ cells were determined as early apoptotic while Annexin V+ 7-AAD+ cells were identified as late apoptotic. Gating strategies used for flow cytometry analyses and sorting are shown in Supplemental Figure 6.

### Immunostaining

To fix cells, 100 µl/500 µl 4% PFA was added to the apical/basolateral chambers and incubated at room temperature for 20 minutes. Both chambers were PBS washed thrice, after which Transwells could be stored in PBS and sealed with parafilm/cling wrap to store at 4°C for future use, or immediately excised for staining.

In some experiments, cells were cytospun to observe *S. pyogenes* infection of individual cells. Cells at ALI were treated with 100 µl Accutase and incubated at room temperature for 15-25 minutes. With a P1000, dissociated cells were gently moved to an Eppendorf containing 600 µl FACS buffer (2% FCS in PBS). Cells were spun at 300 x g for 5 minutes, then resuspended in 160 µl cold FACS buffer (2% FCS) and kept on ice. Cells were then counted with a haemocytometer to obtain a desired number of cells (50,000 - 100,000 cells per 100 µl FACS buffer). To prepare for cytospin, cytospin cuvettes were attached to glass slides using a cytospin clip. Cells were then pipetted into the elbow of the cytospin cuvette and slides were placed into the Cytocentrifuge. Slides were spun at 600 rpm for 5 minutes at Medium acceleration. After spinning, the clip was undone, and the cuvette carefully lifted away from the glass slide. The slides were placed in a clean box to dry overnight. On the following day, slides were fixed with 4% PFA.

Transwell membranes were then surgically excised from the Transwell and placed on a glass slide with the apical side facing up. Using a PAP pen, a hydrophobic barrier was drawn around the sample. Samples were then permeabilised with 0.3% Triton-X100 for 15 minutes at room temperature. Afterwards, samples were PBS washed thrice and subject to blocking with 4% normal donkey serum (Sigma-Aldrich) diluted in PBS. Slides were left to incubate at RT for 1 hour. During this time, primary antibodies were diluted in 4% normal donkey serum. Primary antibodies used in this study are as follows: Polyclonal anti-*S. pyogenes* (ThermoFisher, PA1-73059, diluted 1:100), SFTPC antibody (SantaCruz, sc518029, diluted 1:50), Monoclonal anti-Acetylated Tubulin (Sigma-Aldrich, T7451, diluted 1:500), Polyclonal anti-FOXJ1 (R&D Systems, AF3619, diluted 1:250), Monoclonal anti-MUC5AC (Cell Signalling Tech, 61193, diluted 1:500), Monoclonal anti-SCGB1a1/cc10 (R&D Systems, MAB4218, diluted 1:500), and Polyclonal anti-KRT5/CK5 (Biolegend, 905903/4, diluted 1:500). Following incubation, blocking solution was aspirated and primary antibodies were added to the samples. Slides were incubated overnight at 4°C in a humidity chamber. On the following day, slides were brought back to room temperature and PBS washed thrice (3 minutes each). Secondary antibodies (ThermoFisher, #A32790, #A31573, #A31571, #A10037, #A78948 and/or #A48272) (1:1000) + DAPI (1:10,000) (Sigma-Aldrich, D9542) were diluted in PBS and added to the samples. Slides were incubated at room temperature for 1 hour, protected from light. After incubation, samples were PBS washed thrice (3 minutes each) and then mounted in Fluoromount-G (ThermoFisher, 00-4958-02). The slides were left at room temperature for 4-5 hours (or overnight), protected from light, to allow mounting media to properly set. Slides were stored long-term at −30°C.

Slides were imaged using a Zeiss LSM900 confocal microscope. Images were processed using ImageJ and FIJI.

### Transmission electron microscopy

iAT2s or iAECs on Transwells underwent primary fixation in 2.5% glutaraldehyde (in 0.1M phosphate buffer) and secondary fixation in 2% osmium tetroxide (in 0.1M phosphate buffer). The cells were processed and embedded in resin to provide a support matrix to enable thin sections of the cells to be cut. Excess resin was trimmed off, 0.5 μm semi-thin sections were then cut and stained with methylene blue to provide a good overview under the light microscope. A diamond knife was used to cut the thin sections (50-90 nm). The microscope used was a JEM-1400 TEM (JEOL Ltd.) operating at 80 kV. Images were taken using a 14mp NanoSprint AMT camera and the native AMT software.

### RNA sequencing

RNA was isolated using the ISOLATE II RNA Mini Kit (Bioline, BIO-52073), including on-column DNAse treatment. Total RNA with Ribo-Zero Plus libraries were prepared, then pooled libraries were sequenced using a NovaSeq X (150 bp paired end) (Illumina). Data quality was assessed using FastQC. Reads were aligned separately to the human reference genome (GRCh38) and *S. pyogenes* reference genome (NZ_CP060269.1). All analyses (human or *S. pyogenes*) were therefore performed separately. Counts per gene were determined using featureCounts (GCF_028370295.1 used for *S. pyogenes* gene annotations). EdgeR was used to import and filter the counts. Raw counts were filtered to retain genes with sufficient expression (≥10 counts (human) or ≥1 count (*S. pyogenes*) in at least 2 samples) and duplicate gene names were aggregated by summing counts. Normalisation factors were calculated using the trimmed mean of M-values method in edgeR. A factorial design matrix incorporating cell type, time point, and their interaction was fitted using limma-voom, which applies variance stabilisation and precision weights to RNA-seq count data. Differential expression testing employed linear modelling with empirical Bayes moderation, and p-values were adjusted using the Benjamini-Hochberg method with FDR < 0.05 as the significance threshold.

For cross-cell-type comparisons (human gene expression), interaction contrasts (using limma) were used to identify genes with differential responses to infection between iAT2 and iAEC cells while controlling for baseline expression differences. Gene set enrichment analysis was performed using fgsea with gene sets from the Molecular Signatures Database including Hallmark, KEGG, and Reactome pathways, using ranked lists based on log fold-change values. Gene regulatory network (GRN) analysis was performed to identify transcription factor (TF) regulatory relationships during *S. pyogenes* infection. Differentially expressed TFs (FDR < 0.05, |log2FC| > 0.5) were identified for each comparison, and co-expression networks were constructed by calculating Pearson correlations between TF expression and the top 500 differentially expressed genes (potential targets) using normalized log-CPM values. TF-target edges with |r| ≥ 0.5 were retained to build networks, which were analysed for hub TFs (highly connected regulators). For bacterial gene expression analysis, Gene Ontology (GO) enrichment was performed using Fisher’s exact test to identify biological processes overrepresented among differentially expressed *S. pyogenes* genes (FDR < 0.05, |log2FC| > 1) during infection of iAEC versus iAT2 cells at 6 and 24 hour timepoints.

Data visualisation included principal component analysis (PCA) to assess sample clustering and batch effects, with ellipses showing confidence intervals for experimental groups. Heatmaps of differentially expressed genes were generated using hierarchical clustering with correlation-based distance metrics, scaled by row to standardise expression values across genes. For pathway-level analysis, heatmaps displayed expression patterns of genes within specific gene sets, ordered by experimental conditions to highlight temporal and cell-type-specific responses. Scatter plots comparing log₂ fold changes between experimental conditions were used to identify genes with consistent or divergent responses across comparisons, with points coloured by statistical significance thresholds. Gene set enrichment analysis results were visualised using dot plots, where dot size represents gene set size and colour indicates normalized enrichment score. Data are deposited at GEO: GSE311710.

### Multiplexed cytokine assay

Basolateral media was analysed using a Bio-Plex 27-plex Assay Kit (Bio-Rad, M500KCAF0Y), as per manufacturer’s instructions. Standards, blanks and controls were run in duplicate, and samples (biological triplicates) were analysed neat in singlicate. A Bio-Plex Pro Wash Station was used for washing. Analysis was performed using a Bio-Plex 200 system. Each analyte’s 5PL standard curve was adjusted for >80% fit, and values under the limit of detection were excluded.

### qRT-PCR

Cells were treated with Accutase, collected, then spun down and resuspended in RLY buffer (Meridian, MER-CSA-05101) + 2-ME (ThermoFisher, 21985-023) before RNA extraction using the ISOLATE II RNA Mini Kit as per the manufacturer’s protocol (Bioline, BIO-52073). Absorbance of purified RNA samples was measured using a NanoDrop Spectrophotometer, and RNA concentration (ng/µl) and purity was assessed using the A260/A280 ratio. Tetro cDNA Synthesis Kit (Meridian, MER-BIO-65043) was used to generate complementary DNA (cDNA). qPCR was run for 45 cycles using custom-designed primers with PowerTrack™ SYBR Green Master Mix (ThermoFisher, A46111). The average Ct value for technical triplicates was calculated and normalized to *ACTB* control (for eukaryotic genes) or *gyrA* control (for *S. pyogenes* genes). Fold change was determined over control cells using 2^ΔΔCt^. Biological replicates, as indicated for each figure legend, were run for statistical analyses. The SYBR primer sequences used are provided in Table 1.

**Table 1.**
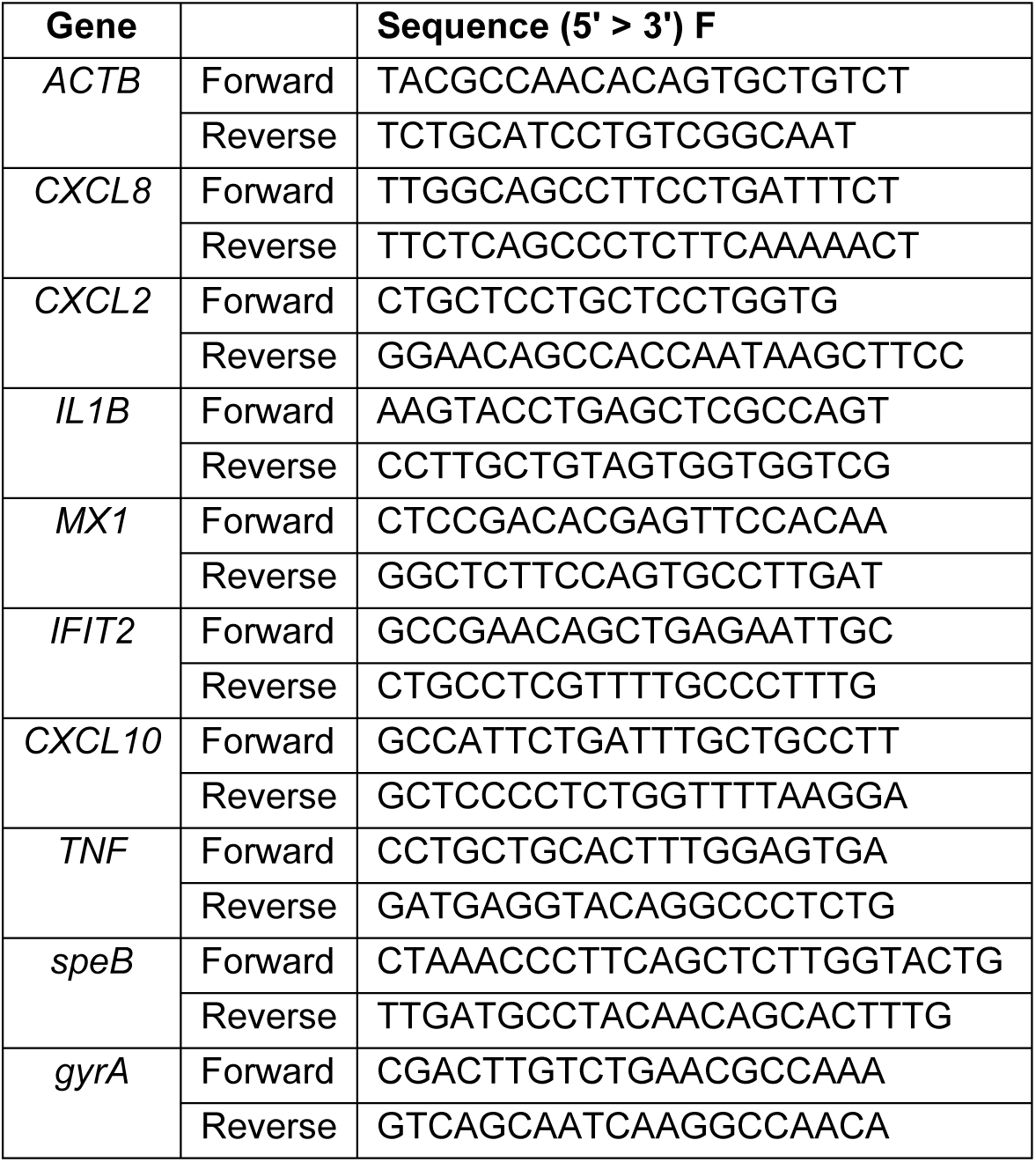
SYBR qRT-PCR primer sequences used in this study.

### Statistics

All datasets were formally assessed for normality using Shapiro-Wilk tests and, where sample size permitted, Kolmogorov-Smirnov tests. Statistical tests are indicated in the figure legend. When comparing two groups an unpaired t-test was used, for three or more groups, a one-way ANOVA (analysis of variance) with Tukey’s multiple comparisons test or a Kruskal-Wallis test with Dunn’s multiple comparisons test (for non-parametric data) was used, and for comparisons with two independent variables (e.g., cell type and time) a two-way ANOVA with Tukey’s multiple comparisons test was used. Significance was determined by a P value of less than 0.05. P value annotations on graphs were as follows: *P < 0.05, **P < 0.01, ***P < 0.001, ****P < 0.0001. All data are represented as mean, with error bars representing standard deviation (SD).

## ACKNOWLEDGMENT

We thank members of the Lung Disease, Immune Development, Blood Disease, Tropical Diseases and Translational Microbiology Groups at the Murdoch Children’s Research Institute (MCRI) for helpful discussions. We are grateful to Meghan McKinnon and Vicki Bennett-Wood for help with TEM preparation and interpretation. We thank Matthew Burton and Eleanor Jones from the MCRI Flow Cytometry and Imaging Facility. This work was supported by the Stafford Fox Medical Research Foundation, L.E.W. Carty Trust, the National Stem Cell Foundation of Australia and the Novo Nordisk Foundation Center for Stem Cell Medicine (grant NNF21CC0073729). The BU3 NGST CRISPRi lines were generated at the Center for Regenerative Medicine, Boston University

## AUTHOR CONTRIBUTIONS

R.B.W. conceptualised the project; J.W. and R.B.W. designed experiments; J.W., D.L.T., S.A., K.P. and R.B.W. performed experiments; J.W., P.V. and R.B.W. performed bioinformatics analyses; M.D., M.R. and F.R. provided support for bioinformatics analyses; H.F., J.V., K.C., K.A. and N.C. grew bacterial stocks, provided support and expertise in *S. pyogenes* infections; C.S., J.O., A.S. and E.S. provided expert input on experimental design and data interpretation; J.W. and R.B.W. wrote the first draft of the manuscript. All authors critically reviewed and approved the final version of the manuscript.

---

**Supplemental Figure 1.** Relates to Figure 2. **(A)** Differentially expressed genes and **(B)** GO enrichment in *S. pyogenes* following infection of iAECs at 24 or 6 hours. **(C)** Differentially expressed genes and **(D)** GO enrichment in *S. pyogenes* following infection of iAT2s at 24 or 6 hours. **(E)** GO enrichment in *S. pyogenes* following infection of iAT2s at 24 hours.

**Supplemental Figure 2.** Relates to Figure 3. **(A-D)** iPSC-derived AECs and AT2s (SCT3010 iPSC line) were matured at air-liquid interface prior to infection with *S. pyogenes* M1_UK_ (1×10^6^ CFU/ml) for 6 or 24 hours. **(A)** *CXCL8*, **(B)** *IL1B*, **(C)** *TNF* and **(D)** *CXCL2* expression were measured by qRT-PCR. **(E-F)** iAT2s were matured at air-liquid interface prior to infection with *S. pyogenes* (M1_UK_) (1×10^6^ CFU/ml) for 24 hours. In some samples the inoculum was removed at 6 hours. *speB* and *CXCL8* were measured by qRT-PCR. **(G-I)** iAECs/iAT2s were matured at air-liquid interface then treated with 1 μg/ml Lipopolysaccharide (gram-negative mimetic) for 24 hours. **(J-L)** iAECs/iAT2s were matured at air-liquid interface then treated with 0.1 or 1 μg/ml Lipoteichoic Acid (gram-positive mimetic) for 24 hours. *CXCL8, CXCL10* and *IL1B* expression was measured by qRT-PCR. **(M-O)** iAECs/iAT2s were infected with *S. pyogenes* M1_UK_ or M75 (1×10^6^ CFU/ml) for 6 hours. IL-6, G-CSF and VEGF levels were measured using a multiplexed cytokine assay. **(P)** iAECs/iAT2s were infected with *S. pyogenes* M1_UK_ (1×10^6^ CFU/ml) for 6 or 24 hours. Viable (Annexin V- 7-AAD-) and early apoptotic cells (Annexin V+ 7-AAD-) were analysed by flow cytometry and normalized to uninfected (UI) cells. **(Q-T)** iPSC-derived type 1 alveolar epithelial cells (iAT1s) were matured at air liquid interface prior to infection with *S. pyogenes* (M1_UK_) (1×10^6^ CFU/ml) for 6 or 24 hours. **(Q)** *speB*, **(R)** *CXCL8,* **(S)** *CXCL2* and **(T)** *IL1B* were measured by qRT-PCR. n = 3 experimental replicates of independent wells of a differentiation; error bars represent SD. Statistical significance was determined using a one way or two way-ANOVA with Tukey’s multiple comparisons test or unpaired t-test; *P < 0.05, **P < 0.01, ***P < 0.001, ****P < 0.0001. UI = uninfected control

**Supplemental Figure 3.** Relates to Figure 4. iPSC-derived AECs were infected with *S. pyogenes* M1_UK_ (1×10^6^ CFU/ml) for 6 or 24 hours. **(A)** Gene set enrichment analysis (GSEA) using the Hallmark database of 6 hours *S. pyogenes* infection in iAECs. **(B)** Gene regulatory network (GRN) of hub transcription factors 24 hours post *S. pyogenes* infection of iAECs, relative to control. **(C)** Basolateral media was analysed using a multiplexed cytokine assay. Heatmap representation of cytokine levels displayed by hierarchical clustering with row scaling (z-score normalised where colour represents relative expression from low (purple) to high (yellow)). **(D)** Absolute levels [pg/mL] of cytokines of interest from (C). n = 3 experimental replicates of independent wells of a differentiation; error bars represent SD. Statistical significance was determined using a one way-ANOVA with Tukey’s multiple comparisons test; *P < 0.05. UI = uninfected control

**Supplemental Figure 4.** Relates to Figure 5. iPSC-derived AT2s were infected with *S. pyogenes* M1_UK_ (1×10^6^ CFU/ml) for 6 or 24 hours. **(A)** Gene set enrichment analysis (GSEA) using the Hallmark database of 6 hours *S. pyogenes* infection in iAT2s. **(B)** GRN of hub transcription factors 24 hours post *S. pyogenes* infection of iAT2s, relative to control. **(C)** Basolateral media was analysed using a multiplexed cytokine assay. Heatmap representation of cytokine levels displayed by hierarchical clustering with row scaling (z-score normalised where colour represents relative expression from low (purple) to high (yellow). Grey values represent samples below the limit of detection). **(D)** Absolute levels [pg/mL] of cytokines of interest from (C). n = 3 experimental replicates of independent wells of a differentiation; error bars represent SD. Statistical significance was determined using a one way-ANOVA with Tukey’s multiple comparisons test; ***P < 0.001, ****P < 0.0001. UI = uninfected control

**Supplemental Figure 5.** Relates to Figure 6. **(A)** Principal component analysis (PCA) of uninfected (control), 6-hour or 24-hour infection iAT2s (red) and iAECs (blue). **(B)** Heatmap showing expression of canonical AT2 and AEC genes. Expression values are row-scaled (z-score normalised) where colour represents relative expression from low (purple) to high (yellow). **(C)** Select DEGs. **(D)** Expression of select IFN receptors. **(E)** Bacterial attachment quantified via CFU assay. **(F)** Expression of *speB.* **(*G*)** Transepithelial electrical resistance (TEER) relative to 0 hours of infection. n = 3 experimental replicates of independent wells of a differentiation; error bars represent SD. Statistical significance was determined using a one way or two way-ANOVA with Tukey’s multiple comparisons test or unpaired t-test

**Supplemental Figure 6.** Summary of flow cytometry analysis and sorting used in this study. **(A)** Endoderm induction was assessed by CXCR4+ cKit+ cells. **(B)** Lung progenitors were sorted based on NKX2-1-GFP expression or **(C)** CD47-hi CD26-lo. **(D)** iPSC-derived basal cells were routinely analysed and/or sorted as necessary for NGFR+ NKX2-1+ or **(E)** NGFR+ EpCAM+. **(F)** iPSC-derived type 2 alveolar epithelial cells were routinely analysed and/or sorted as required for SFTPC+ NKX2-1+ or **(G)** CPM+. **(H)** Apoptosis was assessed based on 7-AAD and Annexin V.

