## Supplemental Figures for "Attenuated interferon signalling in alveolar epithelium limits resistance to *Streptococcus pyogenes*"

### Supplemental Figure 1

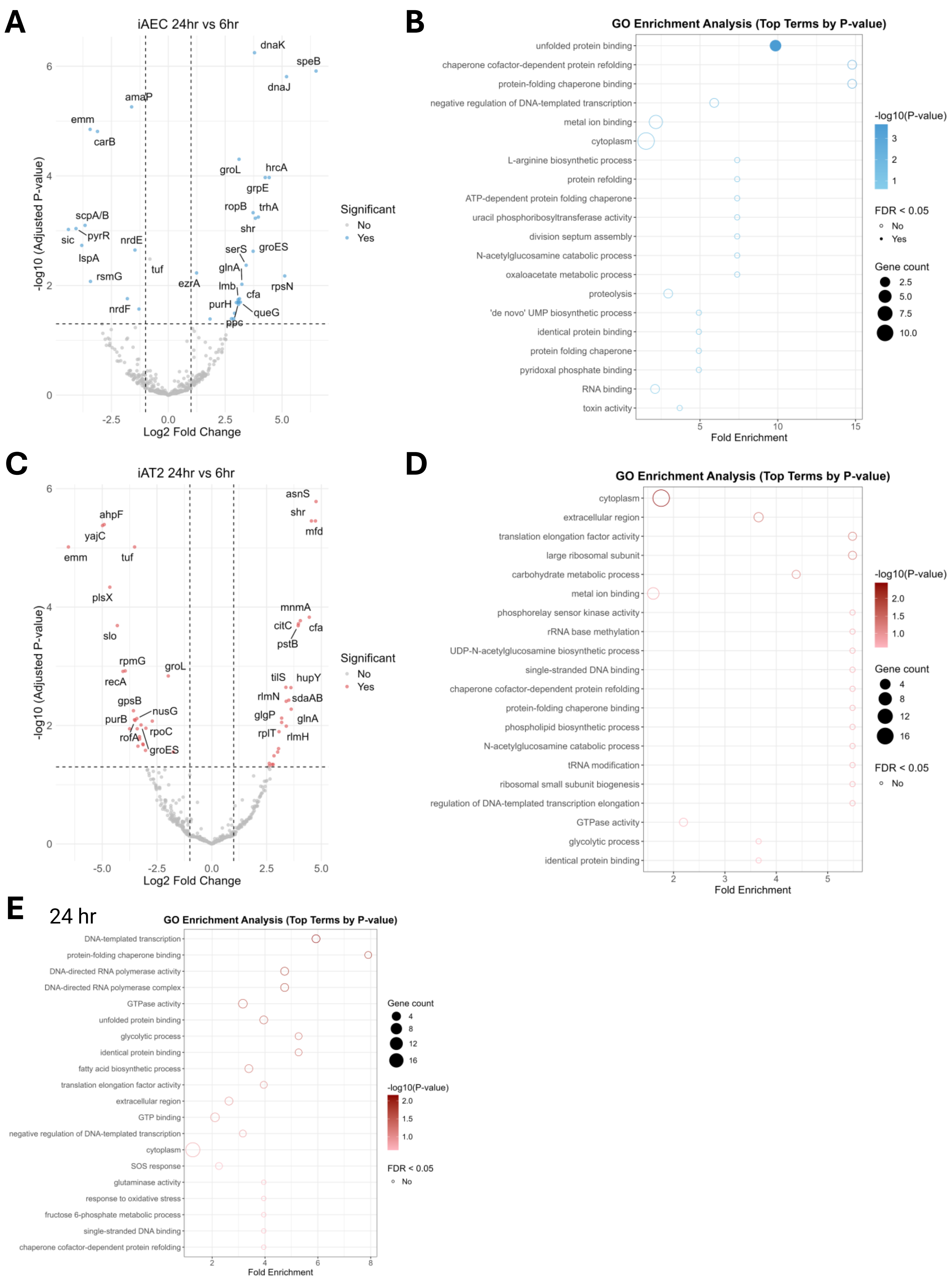

### Supplemental Figure 2

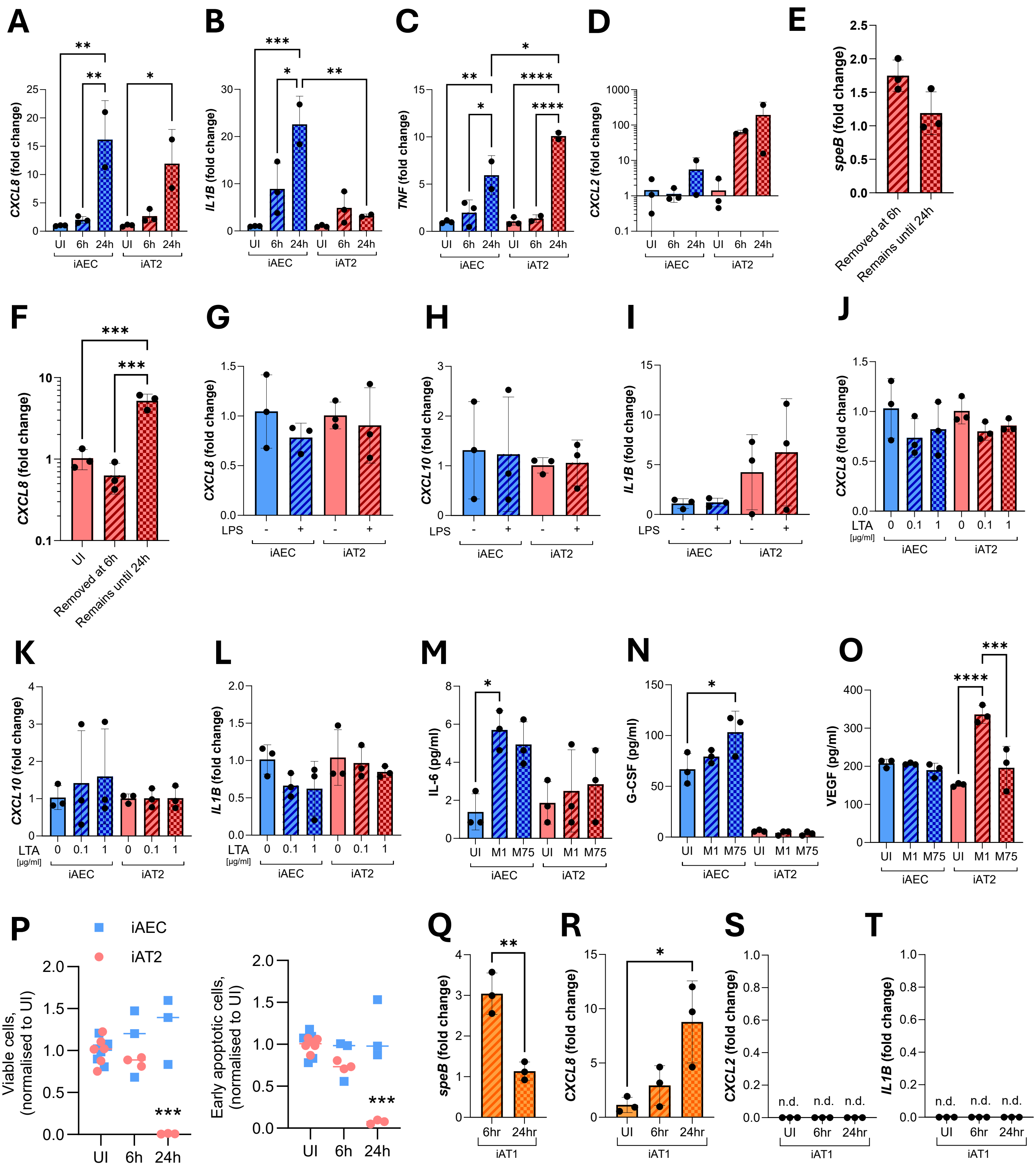

### Supplemental Figure 3

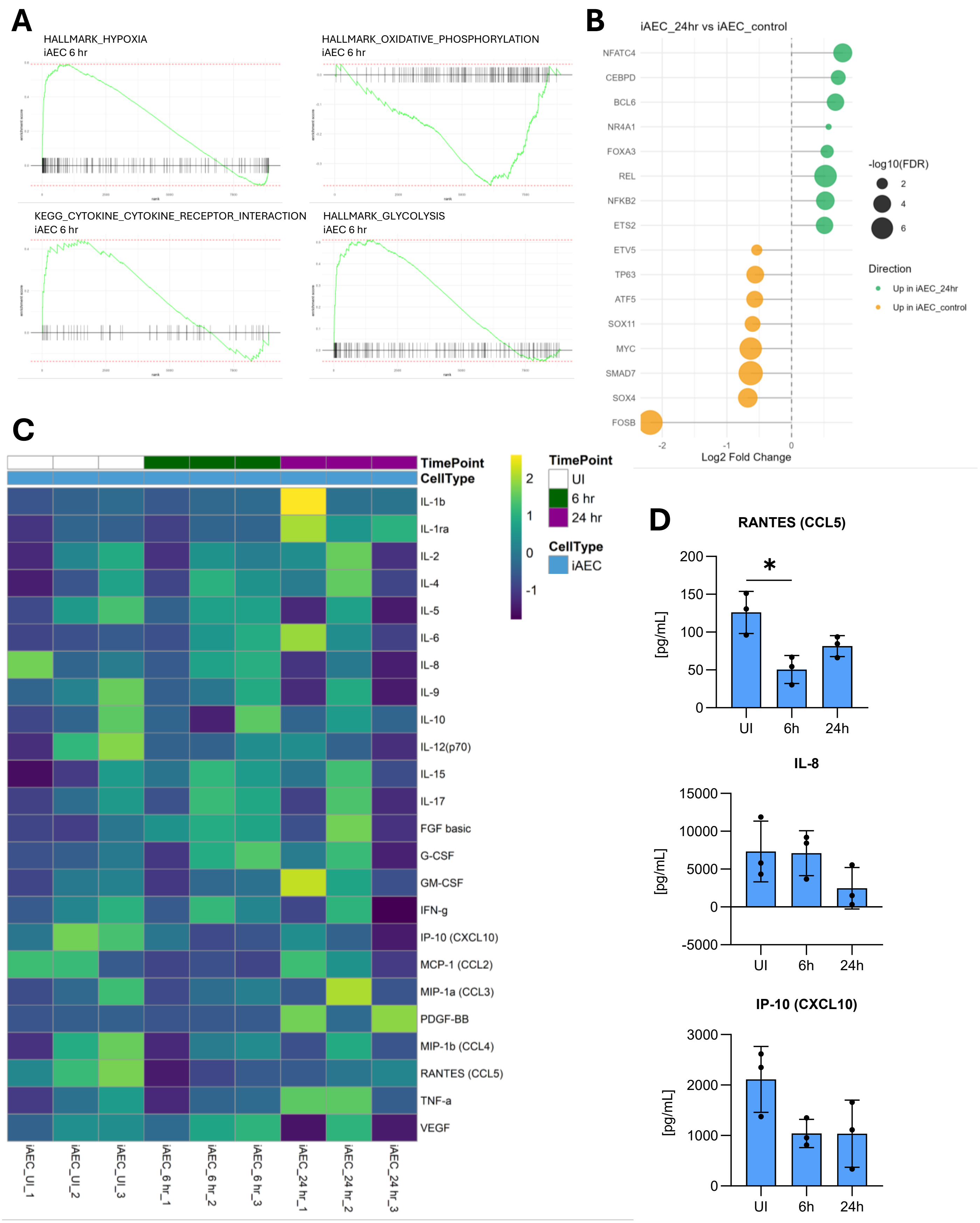

### Supplemental Figure 4

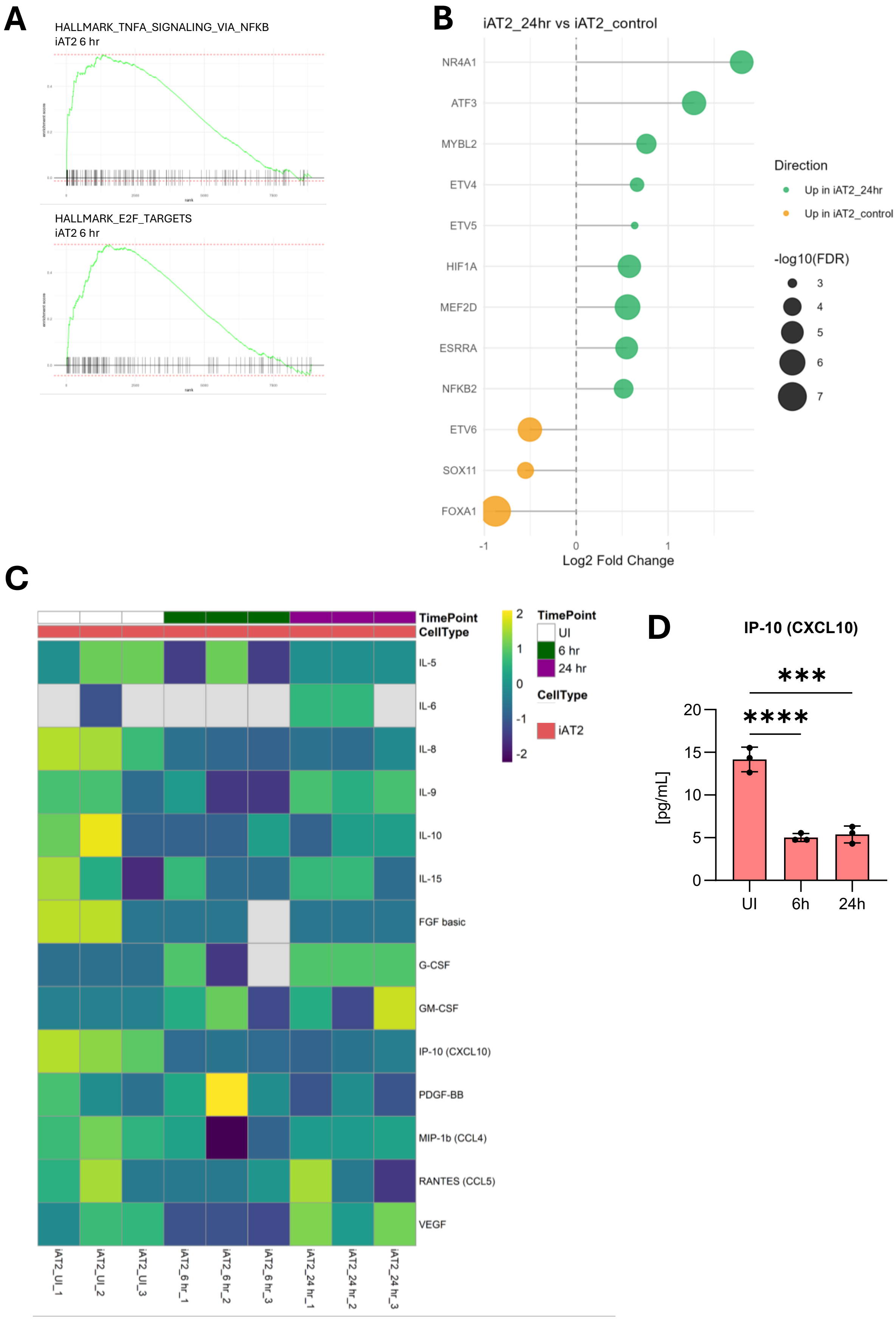

### Supplemental Figure 5

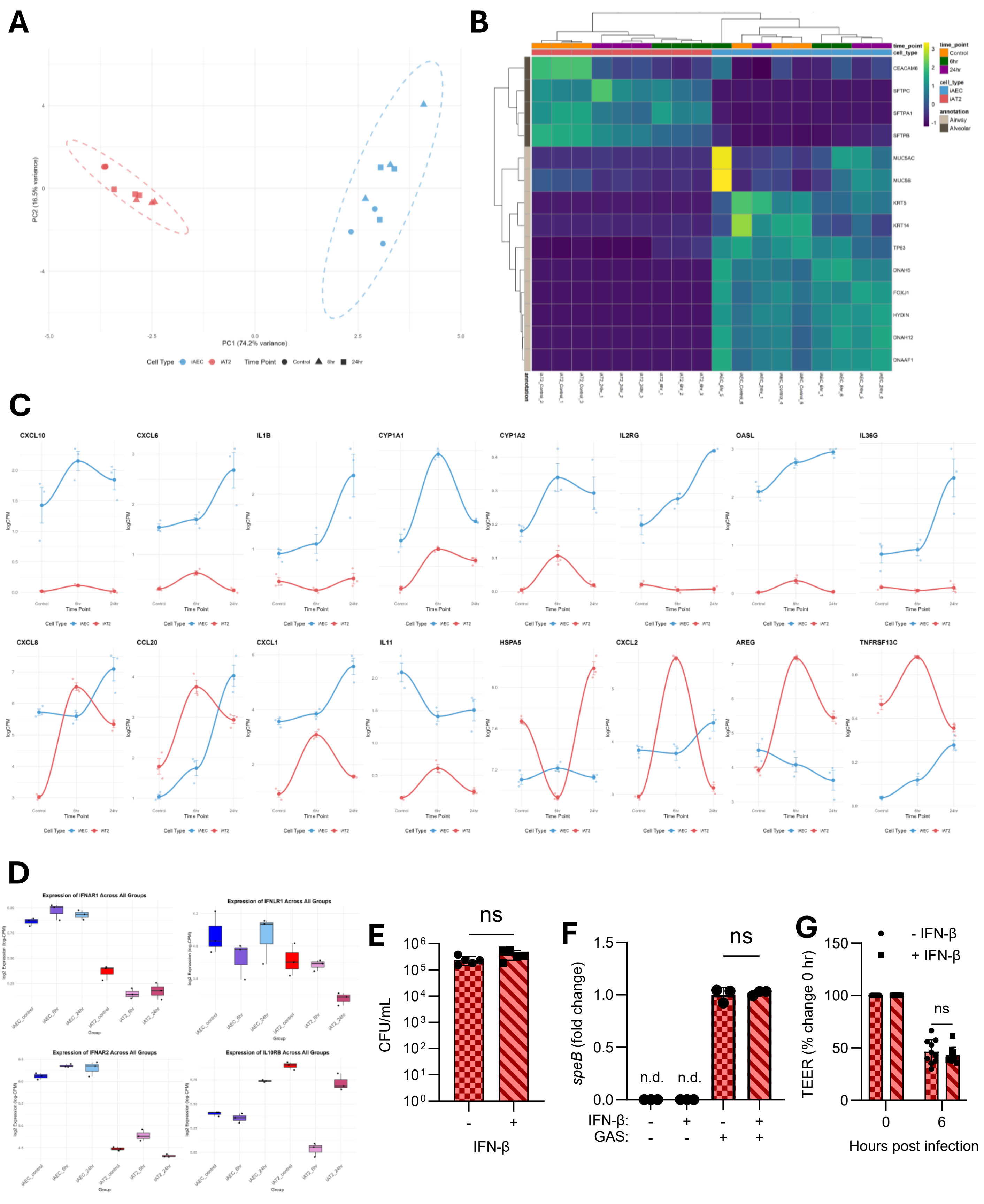

### Supplemental Figure 6

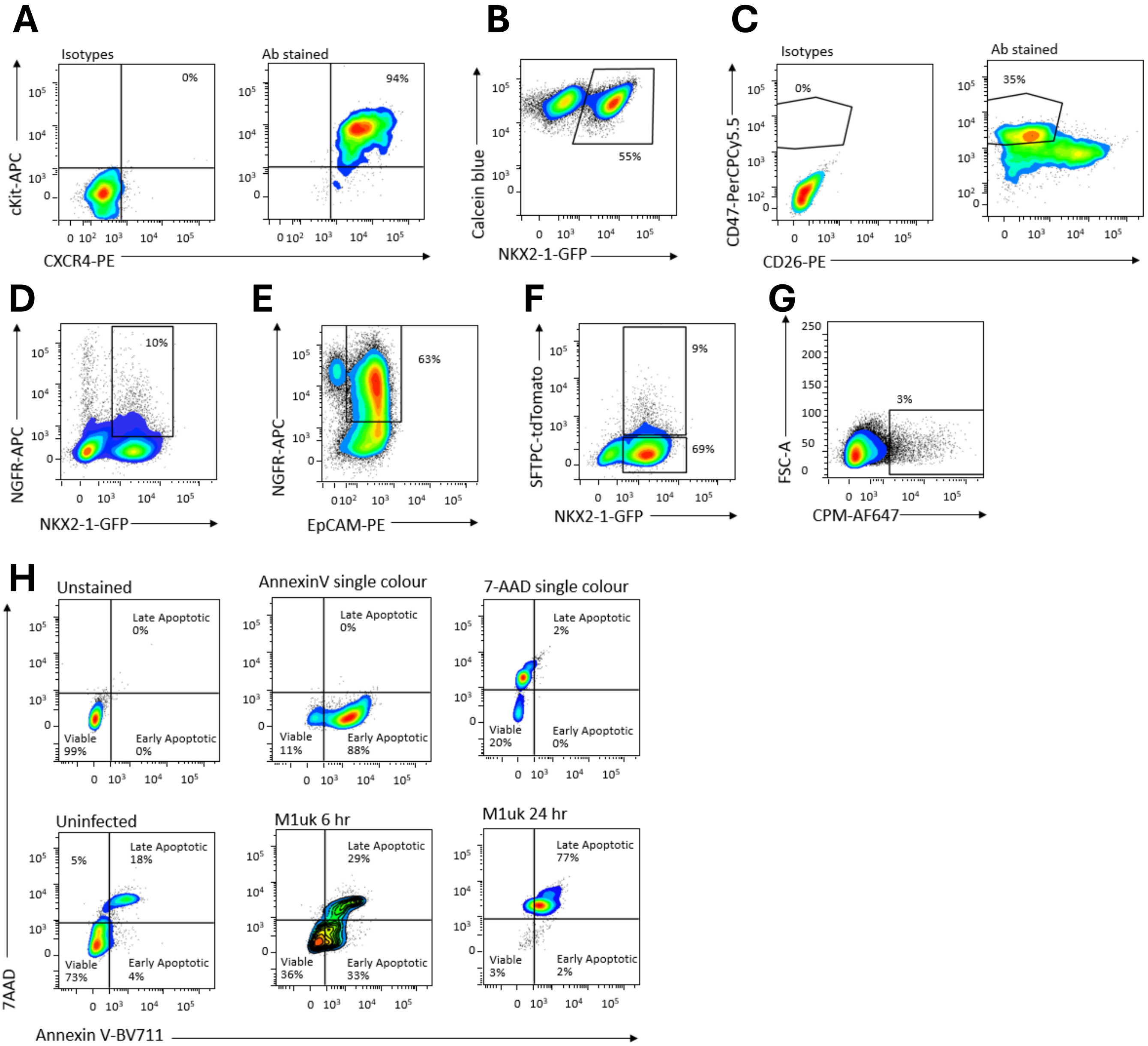
